# Omics Scale Quantitative Mass Spectrometry Imaging of Lipids in Brain Tissue using a Multi-Class Internal Standard Mixture

**DOI:** 10.1101/2023.06.21.546027

**Authors:** Michiel Vandenbosch, Shadrack M. Mutuku, Maria José Q. Mantas, Nathan H. Patterson, Tucker Hallmark, Marc Claesen, Ron M. A. Heeren, Nathan G. Hatcher, Nico Verbeeck, Kim Ekroos, Shane R. Ellis

## Abstract

Mass spectrometry imaging (MSI) has accelerated the understanding of lipid metabolism and spatial distribution in tissues and cells. However, few MSI studies have approached lipid imaging quantitatively and those that have focus on a single lipid class. Herein, we overcome limitation of quantitative MSI (Q-MSI) by using a multi-class internal standard lipid mixture that is sprayed homogenously over the tissue surface with analytical concentrations that reflects endogenous brain lipid levels. Using this approach we have performed Q-MSI for 13 lipid classes representing >200 sum-composition lipid species. This was carried out using both MALDI (negative ion mode) and MALDI-2 (positive ion mode) and pixel-wise normalisation of each lipid species signal to the corresponding class-specific IS an approach analogous to that widely used for shotgun lipidomics from biological extracts. This approach allows pixel concentrations of lipids to be reported in pmol/mm^2^. Q-MSI of lipids covered 3 orders of magnitude in dynamic range and revealed subtle change sin in distribution compared to conventional total-ion-current normalisation approaches. The robustness of the method was evaluated by repeating experiments in two laboratories on biological replicates using both timsTOF and Orbitrap mass spectrometers operated with a ~4-fold difference in mass resolution power. There was a strong overall correlation in the Q-MSI result obtained using the two approaches with outliers mostly rationalised by isobaric interferences that are only resolved with the Orbitrap system or the higher sensitivity of one instrument for particular lipid species, particularly for lipids detected at low intensity. These data provide insight into how mass resolving power can affect Q-MSI data. This approach opens up the possibility of performing large-scale Q-MSI studies across numerous lipid classes and reveal how absolute lipid concentrations vary throughout and between biological tissues.

## Introduction

Mass spectrometry imaging (MSI) is a powerful tool for mapping the spatial distribution lipids^1,2^ and other analytes throughout biological tissues. ^3–5^ Lipids are one of the most common target analytes for MSI, in part due to the relative ease by which several lipid classes are detected and the vital role of lipid metabolism in many biological functions and diseases^6–8^. MSI of lipids has thus been applied in diverse biological applications, including studying solid cancers,^9–11^ neurodegenerative disorders,^12,13^ cardiovascular disease,^14,15^ fatty liver disease,^16^ and lung disease.^17^ Furthermore, recent developments in post-ionization strategies coupled–matrix-assisted laser desorption/ionization (MALDI) such as laser-post-ionization (MALDI-2),^18^ plasma post-ionization^19–21^ and vacuum-ultraviolet-based methods^22^ have significantly expanded the number of lipid classes that can be studied with MSI.^23^

MSI data visualization is based on the mapping of ion intensities across the samples, often following normalisation procedures based on either total-ion current (TIC) or root mean square (RMS) normalisation.^24^ However, ion intensities do not directly correlate to absolute concentrations and can vary significantly depending on the desorption/ionisation method employed, which is not solved through conventional TIC or RMS normalisation. In the case of lipids, ionisation efficiencies are strongly dependent on the lipid class and the presence of other lipid classes, varying by several orders of magnitude across different classes. Regarding imaging, the variation in the chemical composition of species extracted from different anatomical regions within heterogeneous tissue sections plays a crucial role in determining the extent of matrix effects.^25^ As a result, region-specific ionisation efficiencies can depend on the local tissue environment.^26,27^ This can also lead to discrepancies in ion intensities for a given lipid species across a tissue, even if present at equal concentrations.^28,29^ Thus, while MSI has shown much versatility for elucidating region-specific lipid fingerprints and relative changes across tissues, it does not typically allow for absolute quantification (i.e., analyte concentration per unit area or volume within each pixel). Similar challenges in obtaining quantitative data also arise in both shotgun and LC-MS/MS-based lipidomics studies. Currently, these issues are addressed through the use of class-specific stable isotope labelled (SIL) internal standards (IS).^30–32^ These are introduced prior to sample homogenisation and extractions at a single concentration to account for potential lipid losses during sample preparation/analyte recovery and class-specific ionisation behaviours.^33,34^ For a given lipid class, the SIL IS will have an identical ionisation efficiency as endogenous lipids of the same class. Thus, the ratio of endogenous lipid signal to the IS allows for absolute or accurate quantitation of the given lipid class when the IS concentration is known.^35^

Analogous approaches have been adapted for quantitative-MSI (Q-MSI).^36^ In such approaches, a suitable IS (again typically a SIL analogue of the compound of interest) is sprayed evenly across a tissue sample, and the endogenous signal is normalised to the IS signal in each pixel. Q-MSI has most been most widely in the pharmacological context for drug imaging, including imatinib,^37^ clozapine,^38^ and rifampicin,^39^ among others.^40^ In pharmacokinetic/pharmacodynamics (PK/PD) studies, knowing the spatial distribution of pharmaceutical compounds and their localised concentrations is key to a deeper understanding of drug-target engagement, metabolism, biomarker response, changes in the tumour microenvironment and/or resolution of tissue injury.^36^ Standard curves allow for the calibration of signal ratios to absolute concentrations per unit area or volume. However, it is not possible to assess variations in matrix effects across different regions of the analyzed tissue using homogenates. Crucially, Q-MSI has been shown to correlate well with LC-MS/MS data, e.g. the anti-pancreatic cancer drug gemcitabine^41^ and drug candidates in dog liver^42^ using desorption electrospray ionisation (DESI)-MSI and donepezil hydrochloride^43^ in mice brain by MALDI-MSI. Quantitative amounts obtained by Q-MSI often lie within 5% - 20% of those obtained using regional dissections^44^ and LC-MS,^38^ corroborating the validity of the approach.

Q-MSI has been applied using nano-DESI to measure the abundance of phosphatidylcholine (PC) species in specific regions of rat brain tissue sections.^45^ The extraction efficiency of the spray, which depends on the structural properties of tissue, was critical in the estimation of absolute amounts of PC species throughout brain tissue and was corrected for using internal standards.^45^ In another example, Q-MSI of lipids cholesterol was quantified in brain tissue from in a mouse model of Niemann-Pick type 1 disease. ^46^ Q-MSI studies to date has been the targeted nature of the analysis, which means only several targeted analytes or a targeted analyte class (e.g., PC lipids) could be studied, due to the use of a single IS. As MSI is akin to performing a shotgun lipidomics analysis at each pixel, here we have adapted methods from conventional shotgun lipidomics^35^ for quantitation of multiple lipid classes and developed an IS mixture for MSI of brain tissue that contains either SIL, or non-endogenous odd-chain species of 13 lipid classes ubiquitous in brain tissue. Using a combination of MALDI and MALDI-2 as well as orthogonal-time-of-flight and Orbitrap mass analysers we demonstrate the ability to perform Q-MSI on >200 lipid species across positive- and negative-ion mode analyses in a single measurement and investigate the influence of mass resolution on the accuracy of quantitative results.

## Methods

### Chemicals

At the University of Wollongong (UOW) LC-MS hypergrade methanol, chloroform, norharmane and 2,5-dihydroxybenzoic acid (DHB) were purchased from Merck/Sigma-Aldrich (North Ryde NSW, Australia). Haematoxylin and eosin, 1% aqueous were purchased from Point of Care Diagnostics (NSW, Australia). Xylene and quick-hardening mounting medium for microscopy were purchased from Sigma-Aldrich (NSW, Australia). Hematoxylin and Entellan were obtained from Merck (Darmstadt, Germany). Eosin Y was obtained from J.T. Baker (Center Valley, PA, USA) (UM, The Netherlands).

At Maastricht University (UM), water, methanol, and chloroform (all HPLC and ULC/MS grade) were obtained from Biosolve B.V. (Valkenswaard, The Netherlands). Norharmane and 2,5-diydroxybenzoic acid (DHB) were obtained from Sigma Aldrich (St. Louis, MI, USA). Hematoxylin and Entellan were obtained from Merck (Darmstadt, Germany). Eosin Y was obtained from J.T. Baker (Center Valley, PA, USA).

The internal standard mixture (product number #330841 MSI SPLASH) consists of standards for 13 lipid classes developed in collaboration with AVANTI Polar Lipids (Alabaster, Alabama, USA). The composition of the IS mixture used for this work is provided in Table 1.

**Table 1.** Lipid Composition of Internal Standard Mixture used

| Mixture component | Concentration (mg/mL) |
| --- | --- |
| 15:0 – 18:1 (d7) PA | 0.111 |
| 15:0 – 18:1 (d7) PE | 0.100 |
| 15:0 – 18:1 (d7) PG | 0.049 |
| 15:0 – 18:1 (d7) PI | 0.023 |
| 17:0-16:1 PS-d5 | 0.105 |
| 17:0 Lyso PE-d5 | 0.003 |
| C12 Mono-Sulfo Galactosyl(β) Ceramide (d18:1/12:0) | 0.019 |
| C15 Lactosyl(β) Ceramide-d7 (d18:1-d7/15:0) | 0.013 |
| C18 Ceramide-d7 (d18:1/18:0) | 0.011 |
| C17 Glucosyl(β) Ceramide (d18:1/17:0) | 0.133 |
| d18:1 – 18:1 (d9) SM | 0.031 |
| 17:0 Lyso PC-d5 | 0.003 |
| 15:0 - 18:1 (d7) PC | 0.161 |

### Sample Preparation

Fresh frozen brain specimens from wild type (WT) C57BL/6N eight-month aged mice (Jackson Laboratory, ME, USA) were provided by Merck & Co., Inc., Rahway, NJ, USA. Animals were singly-housed, had access to food and water *ad libitum*, and were kept on a 12/12 hour light/dark cycle in a temperature-(22±2°C) and humidity-(approx. 50%) controlled facility. Mice were anesthetized with 3% isoflurane prior to euthanasia, and intact brain specimens were isolated and immediately frozen under dry ice (−80°C) for shipment for analyses at either the University of Wollongong or Maastricht University. At UOW, frozen brain specimens were sectioned at 12-micron (µm) thickness in a cryostat (CM1950 Leica Biosystems, Germany) on pre-cleaned indium tin oxide (ITO)-coated conductive glass slides (Delta Technologies, CO, USA). Slides were kept frozen in a −80°C freezer until analyses. Upon removal, slides were quickly transferred to hygroscopic desiccant (beads) filled boxes and further vacuum-dried further in a desiccation chamber for 20 minutes.

At UM, tissue sectioning was performed on a Leica CM1860 UV (Wetzlar, Germany) at −20°C. 12 µm thick sections were transferred to conductive ITO coated microscope glass slides (Delta Technologies, Loveland, MN, USA). Slides were stored in a −80°C freezer until analyses and vacuum dried as described above. All animal studies were performed under the approval of the Merck & Co., Inc., Rahway, NJ, USA, Institutional Animal Care and Use Committee and endorsed by the Animal Ethics Committee of the University of Wollongong. For the research conducted at Maastricht University, an exemption was granted for an ethics application and the study received approval.

### Internal Standard and Matrix Application

At both institutes, the internal standard mixture (IS mix) was diluted ten-fold in LC-MS grade methanol prior to deposition onto tissue sections. An off-line 2.5 mL Leur lock gas-tight syringe (Trajan Scientific, Victoria, Australia) in combination with a syringe pump was connected to the TM-Sprayer (HTX Technologies, USA) for spray coating of the MALDI SPLASH mix. The settings for IS mix deposition were: temperature, 50°C; number of passes, 16 layers; flow rate, 0.06 μL/min; velocity, 1200 mm/min; track spacing, 2 mm; gas flow rate, 2 L/min and drying time in between passes, 30 sec. Identical spraying parameters were used across both labs in Wollongong and Maastricht. The amount of IS deposited in µg/mm^2^ was calculated by multiplying the analyte concentration (µg/mL), flow rate (mL/min), time (min), surface area sprayed (mm^2^) and number of passes (layers). This value was then divided by the average molecular weight (Da) and the dilution factor, yielding concentration per unit area expressed as picomoles per mm^2^ (pmol/mm^2^). The resulting concentrations for each IS are provided in *Supporting Information Table S1*. Slides were then immediately coated with MALDI matrix. Norharmane and DHB matrices were used for negative and positive ion mode imaging, respectively, and were both dissolved in chloroform-methanol mixture (2:1 v/v). Matrix was deposited on tissue samples using the same TM-Sprayer (HTX Technologies, USA). For negative ion mode analyses, the spray settings were matrix concentration, 7 mg/mL norharmane; temperature, 30°C; number of passes, 15 layers; flow rate, 120 μL/min; velocity, 1200 mm/min; track spacing, 3 mm; gas flow rate, 10 psi and time in between passes, 30 sec. For positive ion mode analyses, the spray parameters were: matrix concentration, 15 mg/mL DHB; temperature, 30°C; number of passes, 10 layers; flow rate, 120 μL/min; velocity, 1200 mm/min; track spacing, 3 mm; gas flow rate, 2 L/min and time in between passes, 30 sec.

### Mass Spectrometry Imaging

#### Orbitrap Elite

At the UOW, tissue sections were analysed using an Orbitrap Elite mass spectrometer (Thermo Fisher Scientific GmbH, Bremen, Germany) equipped with a dual MALDI/ESI Injector (Spectroglyph LLC, Kennewick, WA, USA). All acquisitions were conducted at a pixel size of 75 μm (x, y), MALDI laser pulse energy of 1.0 µJ/pulse (measured after an external attenuator), an injection time of 250 ms, automatic gain control turned off and a mass resolution of 240,000 @ *m/z* 400. Negative ion mode analysis was conducted using conventional MALDI (i.e. without MALDI-2) using laser repetition rate of 500 Hz and a mass range of *m/z* 180-2,000. Positive-ion mode analysis was conducted using MALDI-2 and a mass range of *m/z* 350-2,000. Laser post-ionisation was achieved using a 266 nm laser (NanoDPSS, Litron lasers, Rugby, UK) operating with 500 µJ pulses entering the ion source. Both lasers were operated at 300 Hz and an inter-pulse delay of 20 µs. Further details on the MALDI-2 setup can be found in reference.^47^ For both Orbitrap and timsTOF Flex data was acquired from three biological replicates sent to each lab (i.e. UOW and UM; 6 mouse brain samples in total).

#### MALDI-2-timsTOF Flex

At UM, tissue sections were imaged on a MALDI-2 timsTOF flex (Bruker Daltonik, Bremen, Germany). The mass resolution of this instrument is calculated to be circa 54,000at *m/z* 700. Primary ionization of the material was achieved with a SmartBeam 3D 355 nm laser. Data was acquired at a pixel size of 30 μm (x, y) using a beam scan area of 26×26 µm and a mass range of *m/z* 300-1000. For negative mode measurements, the laser was operated at 10 kHz with 200 shots accumulated per pixel and data acquired across a mass range of *m/z* 300-1000. For positive mode measurements using MALDI-2, both lasers were operated at 1 kHz with 50 laser shots accumulated per pixel and data acquired across a mass range of *m/z* 350-1,000. The MALDI-2 laser (NL204-1K-FH, Ekspla, Lithuania) was operated at 500 µJ/pulse with a pulse delay time of 10 µs. The instrument was calibrated using red phosphorus prior to each measurement. In both cases glass slides after MSI were subjected to haematoxylin and eosin (H&E) staining method (see protocol in Supporting Information).

### Data Processing and Analyses

#### Orbitrap Elite Lipid Identification Target lists

Orbitrap raw data was internally recalibrated using Recal Offline software. Masses used for recalibration are provided in *Supporting Information Table S2*. MSI files were then converted to mzML using Proteowizard msConvert GUI^48^ before conversion to imzML using Lipostar MSI software (Molecular Horizon Srl, Perguia, Italy).^49^ Data was imported into Lipostar MSI to generate initial lipid target lists. Import parameters were: Savitzky-Golay smoothing was performed at window size, 7 points; degree, 2; iterations, 1; for peak picking the minimum SNR was set at 0.00; noise window size, 0.10 amu; minimum absolute intensity at 0.00. Peaks belw 0.50% and 0.20% of the base peak were discarded, for negative mode and positive mode, respectively. Dataset *m/z* tolerance was set at ±5.00 ppm; minimum peak frequency, 2.00%; spatial chaos, 0.7; isotopic clustering abundance deviation, 30%; and *m/z* image correlation threshold, 0.50. Using the resulting peak list initial lipid identifications were achieved using the identification functions in LipostarMSI. A single peak list derived from a merged dataset was used for the identification tool and tentative MS1 level lipid annotation are based on the LMSD “bulk” structures database ^50,51^. Identified peaks were filtered to a 3 ppm tolerance, non-identified compounds were removed and “approved” as even chain. The resulting ID compounds were further filtered according to lipid class and exported as .csv files. The matched ID compounds were manually checked for isobaric lipid sum composition and peak lists were separated according to adduct types. Identification lists for each lipid class were manually curated and then cross-checked against a recently published in-depth brain lipidomics study.^52^ Lipid species that were not detected in both studies, were removed. The full list of lipids is provided in Supporting Information.

### timsTOF data processing

timsTOF data was internally recalibrated using DataAnalysis software (Bruker, Bremen, Germany). A linear lock mass recalibration was applied using the masses as provided in *Supporting Information Table S2*. Subsequently, MALDI-MSI data was imported and analyzed in SCiLS lab software (Bruker, Bremen, Germany) where MSI files were converted to imzML for further analysis.

## Data Analysis and Visualisation

Intensities of the various species of interest, i.e. the target lipid species and internal standards, were extracted through binning of the spectra around the computed corresponding *m/z* value. For this, a window of 12 and 3 ppm was set for the timsTOF and Orbitrap data, respectively. The maximum intensity per pixel within these windows was taken as the ion intensity for the corresponding pixel. Images were normalized by conducting a pixel-wise division of the original *m/z* image by the reference *m/z* image (internal standard). The resultant scaled pixel values were multiplied by the corresponding internal standard concentration to produce an absolute concentration image for each lipid class. Pixels with missing values, resulting from zero-division, were replaced by the median value of a surrounding 3×3 pixel window. Winsorizing was employed to eliminate hotspots in the images, with the .99, .75, and .99 quantiles for the original m/z image, reference m/z image, and normalized image, respectively. Winsorizing is a robust method for outlier removal where pixel intensities above a specified quantile are replaced with the value of that quantile.^53^ The reference *m/z* images used a different quantile due to the pronounced intensity of the internal standard in the matrix surrounding the tissue. For each MSI experiment, a digital 10x/20x H&E stained whole slide microscopy image of a post-MALDI imaging tissue section was acquired. The web-based digital pathology tool, Annotation Studio (Aspect Analytics NV) was used to annotate brain regions in the images with one of five labels: Prefrontal Cortex/Isocortex, Midbrain, Hindbrain, Basal Ganglia, and Cerebellum.

The MSI and neighbouring microscopy images were co-registered using a proprietary landmark-based, non-rigid registration pipeline (Aspect Analytics NV). After registration, the identified anatomical regions of interest (ROIs) were mapped onto the MSI data, facilitating the extraction of ion intensities from the pixels within these ROIs. This enabled the calculation of mean concentrations for each anatomical region across tissue sections

## Results and Discussion

### Optimisation of IS mix Concentration

A key requirement for the IS mix is that the signal for each lipid standard is reflective of the signals obtained for endogenous lipids of the same class. As the abundance of different lipid classes varies substantially due to both differing physiological and biochemical concentrations and class-specific ionisation efficiencies, the concentration of each IS component must be fine-tuned. The final composition and concentration of the IS mix, shown in Table 1, was determined following an iterative process using both the timsTOF Flex and Orbitrap Elite, which tested for possible interferences with the expected IS peaks. **Figure 1a** shows extracted negative-ion mode mass spectra of endogenous lipid species for LPE, PE, PG, PI, PS and SHexCer species their respective reference internal standard (marked in red) after averaging all on-tissue pixels from one brain section measured using the Orbitrap Elite. For comparative data acquired from the timsTOF Flex and for other lipid classes see *Supporting Information Figures S1 and S2*. As intended, the IS reference peaks for LPE (*m/z* 471.3253), PE (*m/z* 709.5519), PI (*m/z* 828.5625), PS (*m/z* 751.5291) and SHexCer (*m/z* 722.4519) have abundances similar to moderately abundant endogenous lipid species of each class. Given the low signals of PG species and in order to ensure detectability of the IS signal in every pixel, the concentration of the PG IS (*m/z* 740.5464) was intentionally designed to supersede the most intense endogenous peak.

**Figure 1.**
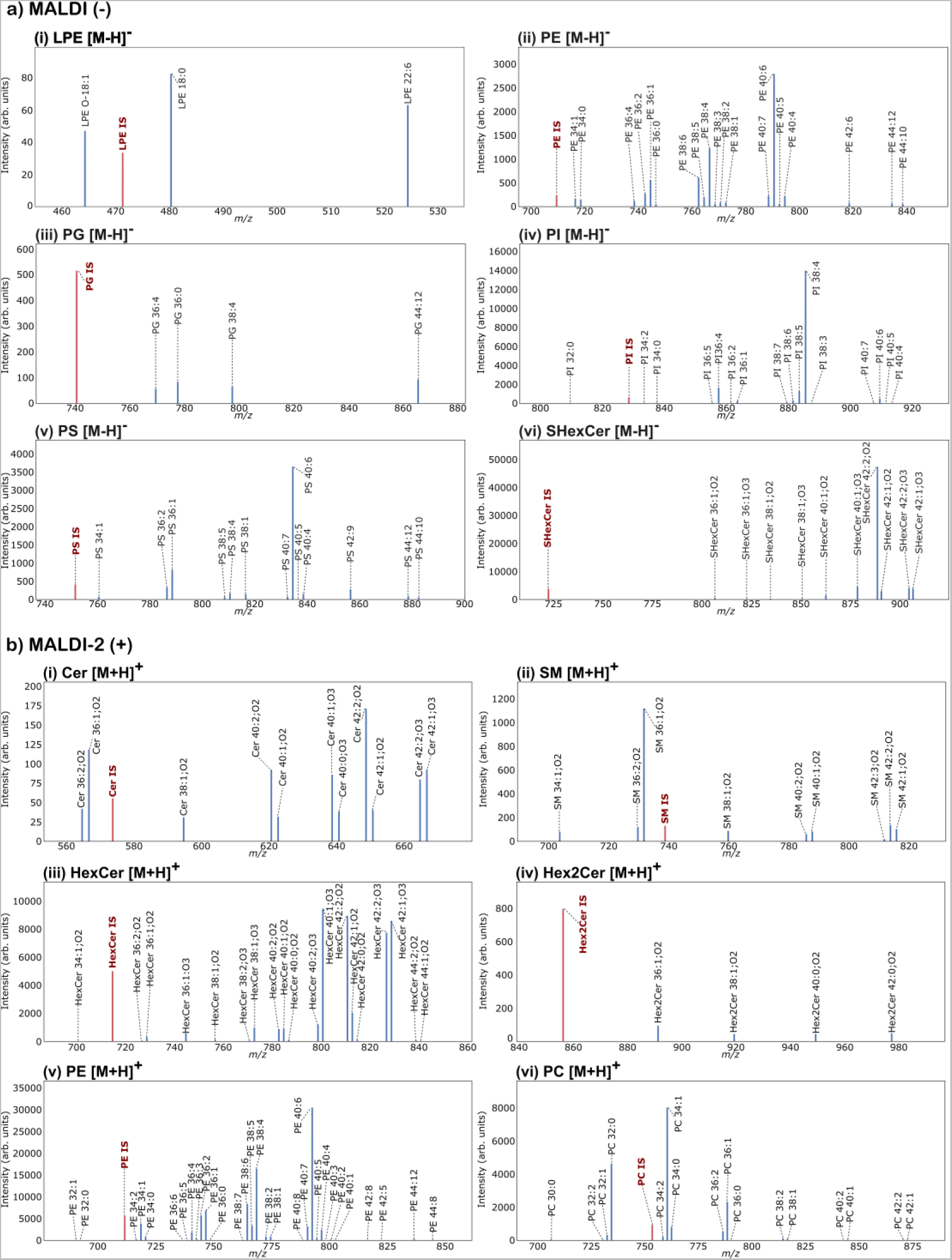
Extracted Orbitrap mass spectra averaged over entire mouse brain tissue region. (a) Peaks corresponding to (i) LPE, (ii) PE, (iii) PG, (iv) PI, (v) PS and (vi) SHexCer lipid species detected as [M-H]^−^ ions in negative ion mode. (b) Peaks corresponding to (i) Cer, (ii) SM, (iii) HexCer, (iv) Hex2Cer, (v) PE and (vi) PC lipid species detected as [M+H]^+^ ions in positive ion mode. Reference IS peak shown in red and endogenous lipid species shown in blue. Similar data for other lipid classes and those acquired using the timsTOF Flex is provided in the *Supporting Information Figure S2*.

**Figure 1b** shows the corresponding positive-ion mode data acquired using MALDI-2 with IS reference observed for the [M+H]^+^ ions of Cer (*m/z* 573.5946), SM (*m/z* 738.6470), HexCer (*m/z* 714.578), Hex2Cer (*m/z* 855.6533), PE (*m/z* 711.5664) and PC (*m/z* 753.6134). Similar to the PG IS, the higher abundance of the IS for Hex2Cer was deliberately chosen to ensure consistent detection of the IS across the tissue. These data demonstrate the suitability of the chosen concentrations within the IS mix for lipid imaging of brain tissue.

### Application of Multi-Class IS Mix to Lipid MSI

Next, we evaluated the effect of internal standard (IS) normalisation on lipid imaging. The *m/z* images were plotted with a ±3 ppm theoretical mass window of the chosen lipid species for negative ion (Figure 2a) and positive-ion mode (Figure 2b) for the Orbitrap Elite data. For each representative lipid species the original ion image is shown on the left panel, the IS distributions in the centre and the IS normalised lipid distribution is shown on the right. The reference images show mostly an intense uniform IS distribution on the off-tissue (boundary margin) areas, reflective of the enhanced ionisation efficiency of standards off the tissue surface, and a non-homogenous spatial distribution across the on-tissue areas, reflective of the region-specific ionisation efficiencies that the IS intends to correct. In both negative and positive-ion mode data, the original images are broadly consistent with the IS normalised images, however subtle differences can be observed that highlight the effect of IS normalisation. For example, in negative ion mode analyses, the IS normalised images of PI 38:4 and PS 36:2 ([M-H]^−^ ions) show higher lipid abundance in the brain stem regions compared to the original data (*Supporting Information Figure S3*). Another benefit of IS normalization, is the correction for region-specific distribution of endogenous species to reveal regions of higher concentration. For example, SHexCer 42:2;O2 showed patterns of localisation in the white matter (WM) fibre tracts adjacent to the caudoputamen and pallidum inclusive of the cerebellum after IS normalisation. Indeed, this correction is beneficial for many lipids including low abundant species such as SHexCer 36:1;O2, which is more evidently distributed within WM regions (*Figure S4*). Additionally, despite the fact that PI 36:2 seems uniformly distributed across the brain without normalization, it appears more localised within the WM area of the basal ganglia and brain stem, whereas PI 38:6 initially more intense across the cerebellum was higher in the hindbrain after IS normalization (*Supporting Information Figure S4*).

**Figure 2.**
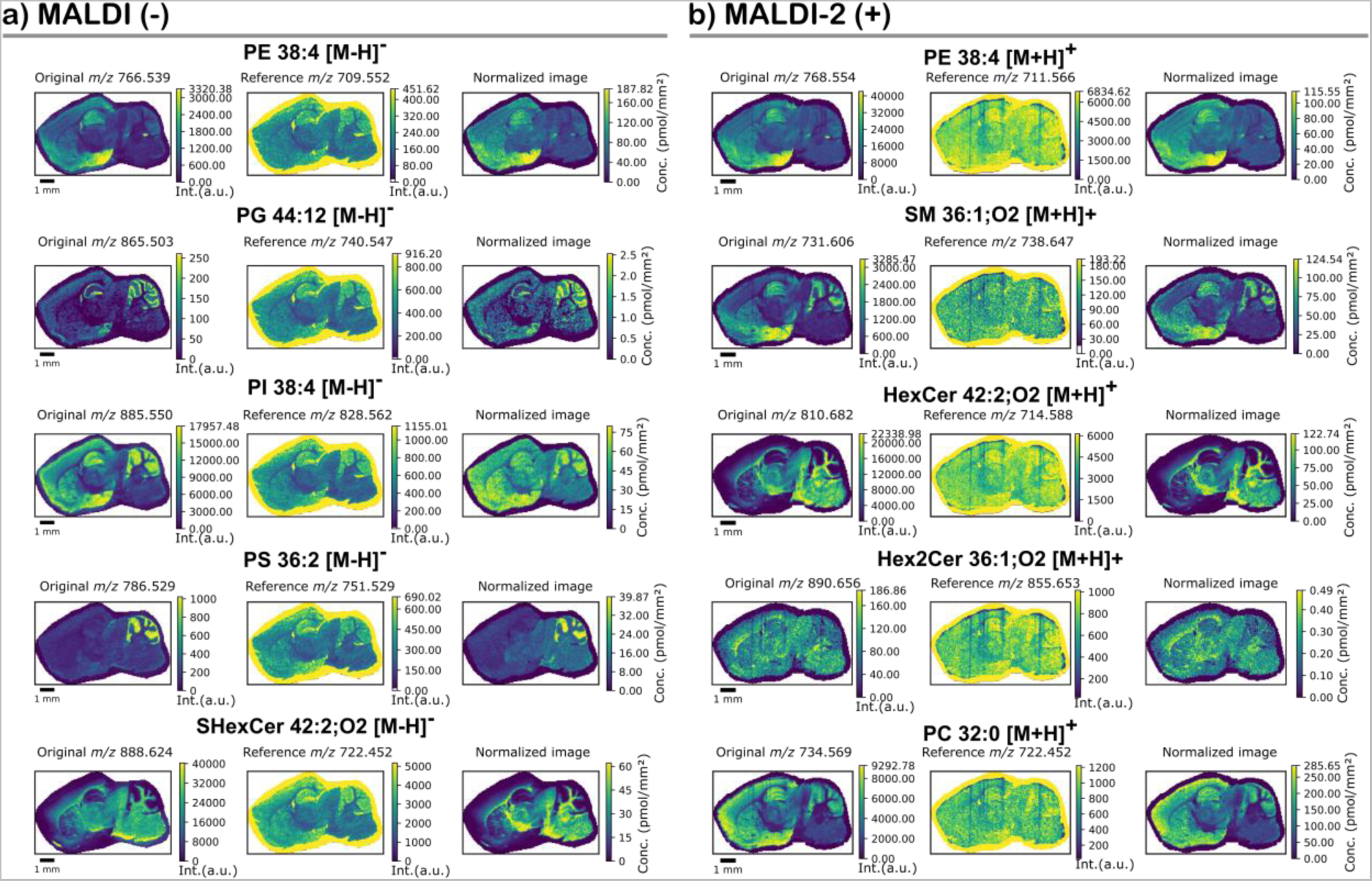
Representative internal standard (IS) normalised ion images for different lipids species detected in (a) negative ion mode using MALDI and (b) positive ion mode using MALDI-2 acquired using the Orbitrap Elite. For each lipid species the original ion image is shown on the left, the class-specific internal standard ion image show in the centre and the IS normalised ion image shown on the right. Intensities for each lipid species were selected using an *m/z* window of ±3.0 ppm compared to the theoretical *m/z* of the lipid species. (A) shows representatives negative-ion mode data using MALDI-MSI for the [M-H]^−^ ions of PE 38:4, PG 44:12^−^, PI 38:4 PS 36:2, and SHexCer 42:2;O. (B). shows representative positive ion mode data using MALDI-2-MSI for the [M+H]^+^ ions of PE 38:4, SM 36:1;O2, HexCer 42:2;O2, Hex2Cer 42:2;O2 and PC 36:0. Similar imaging data from the timsTOF Flex can be found in Figure S6.

Similar results were obtained in positive ion mode with the distributions of selected PC, SM, PE, HexCer and Hex2Cer shown in Figure 2b. Analogous to Figure 2a, the original (left-hand panels) and IS normalised (right-hand panels) images show the same general distributions, but with the benefit of being able to quantify lipid concentrations (see below). Another notable benefit of IS normalisation is the ability to correct for strips or streaks in ion images that is sometimes observed using MALDI-2 (Figure 2b). This can be caused by partial blockage of the MALDI-2 laser beam for example by a large matrix crystal, slight laser alignment drift or topographical changes on tissue edges. This effect is particularly visible in the images of the [M+H]^+^ ions of the PE and HexCer ISs as vertical lines (i.e. the direction of stage raster). Since the same effect is seen on endogenous species as on the IS, this is corrected after IS normalization. In all, reproducible results were obtained across biological replicates, with several examples shown in *Figure S5*. Comparative data acquired from the timsTOF Flex is provided in the *Supporting Information Figure S6*.

### Region-Specific Quantitation of Lipids using Q-MSI

Next, we deployed our multiplexed Q-MSI approach to reveal lipid concentrations within histologically-defined brain regions. Brain regions were defined on the H&E-stained tissue sections which were then co-registered with MSI data (see methods). **Figure 3a** shows the concentrations (pmol/mm^2^) in negative ion mode for PE, PS, PI and SHexCer as [M-H]^−^ acquired using the Orbitrap Elite. Lipid concentrations across the presented classes had concentrations of defined sum-composition species varying from several hundred pmol/mm^2^ to less than 1 pmol/mm^2^. These data also highlight the effect of class specific ionisation biases. For example, the base peak in negative-ion mode is typically either PI 38:4 or SHexCer 42:2;O2 depending on the brain region, however several PE and PS species have absolute concentrations several fold higher than both PI 38:4 or SHexCer 42:2;O2. These results are also consistent with those reported by Fitzner *et al* which also showed higher amounts of PE and PS in brain tissue compared to PI and SHexCer.^52^

**Figure 3.**
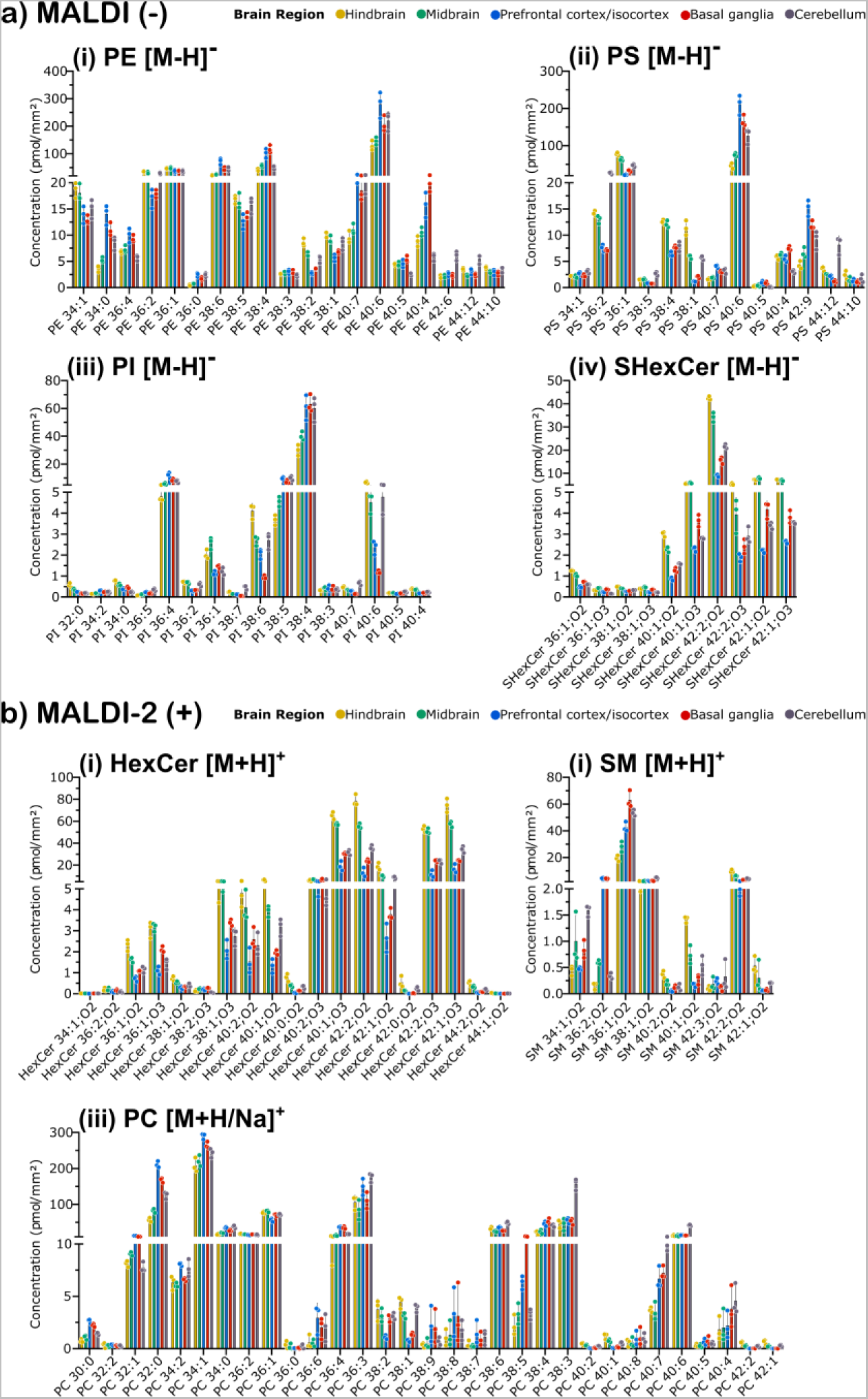
Mean concentration of lipid species from quantitative mass spectrometry imaging in five different brain regions acquired using the Orbitrap Elite. Regions of interest (ROI) from sagittal brain tissue sections are colour-coded hindbrain – orange, midbrain – green, prefrontal cortex/isocortex – blue, basal ganglia – red, and cerebellum – grey. **(a)** Selected lipid classes detected by regular MALDI-MSI negative ion mode lipid species measured as [M-H]^−^ ions (i) PE, (ii) PS, (iii) PI, and (iv) SHexCer. **(b)** Selected lipid classes detected by regular MALDI-2 MSI positive ion mode (i) HexCer, (ii) SM lipid species measured as [M+H]^+^ ions, and (iii) a combined non-isobaric list of PC [M+H]^+^ and PC [M+Na]^+^ adducts. Each bar represent the average concentration from n=3 biological replicates and error bars represent ± 1 standard deviation. Individual data points from each replicate are provided for each species.

**Figures 3b** shows the Q-MSI obtained for all detected lipid species in positive ion mode using the Orbitrap Elite across SM, HexCer as [M+H]^+^ ions, and PC. For each PC species, individual adducts were chosen to avoid isobaric interference between sodiated and protonated PC lipids. Generally this meant that PC species with few double bonds were analysed as the protonated species and more unsaturated species were analysed as the sodiated species *(Supporting Information Table S3)*. This approach was used given that the [M+K]^+^ adduct of the PC IS could not be resolved from the [M+1] isotope of protonated PS 36:1. As expected PC species yielded the highest concentrations with several species having concentrations >100 pmol/mm^2^ other myelin-specific lipids such as HexCer yielded concentrations approaching 80 pmol/mm^2^ for the most abundant HexCer42:2;O2. Region-specific concentrations of additional lipid classes and for data acquired using the timsTOF are provided in the *Supporting Information Figures S7, S8 and S9*.

To validate our quantitative technique we compared spatial lipidomics data from this study to other bulk lipidomics studies that reported lipid concentrations in units our data could be compared. After accounting for section thickness and assuming a tissue density of 1 g/cm^3^, we have converted our values to match prior literature to allow for direct comparisons. Eiersbrock *et al*.,^54^ reported concentrations of PC 34:1 of 9.07±0.9 nmol/mm^3^ and 7.0±1.3 nmol/mm^3^ for the WM and the molecular layer (ML), respectively, while Choi *et* al.,^55^ reported a concentration of 10.06 nmol/mm^3^ for whole mouse brain tissue. Averaged across the entire tissue, we obtained a concentration of PC 34:1 as 16.17 nmol/mm^3^ after accounting for tissue thickness, with similar consistency also found for PC 38:4 and PC 40:6 at 2.49 nmol/mm^3^ and 1.06 nmol/mm^3^ respectively. Both studies also reported concentrations of PE 34:1 between 0.46 and 1.46 nmol/mm^3^ depending on brain region, which again compares favourably with our data of 0.97 nmol/mm^3^ Eiersbrock *et al* also reported a concentration of 10.2±1.4 and 0.37±0.181 nmol/mm^3^ for SHexCer 42:2;O2 for the WM and ML, respectively, which is consistent with the value we obtained of 1.47 nmol/ mm^3^ (note higher values are observed in the WM regions, consistent with Eiersbrock *et* al). Our results for PC quantitation are also generally consistent with reported values by Jadoul *et al*.,^56^ for PC 34:1, PC 36:1 and PC 32:0 protonated species: 12,282 versus 14,980 µg/g, 3,682 versus 6,347 µg/g and 6,527 versus 4,594 µg/g, respectively, who used a spiked tissue homogenate and an isotopically labelled PC IS to quantify PC species in mouse brain tissue. In addition, we have compared our absolute ratio of PC to PE lipids to data acquired using LC-MS with excellent agreement found between the methods (*Supporting Information Figure S10*). Taken together, these data provide confidence that absolute concentrations reported by our multiplexed Q-MSI approach (after averaging across tissue regions for comparison) yield similar results acquired following lipid extraction quantitation using LC-MS or shotgun lipidomics.

### Multiplatform Comparison on Multiplexed Q-MSI Approach

Overall, similar quantitative results and ion images were observed between the Orbitrap and timsTOF data (*Supporting Information Figures S6, S7, S8 and S9*). The timsTOF yielded superior image quality for IS normalised ceramide and LPE species with several examples highlighting this shown in **Figure 4**. This is largely due to better detection of the respective LPE and Cer IS species using the timsTOF under the employed conditions - which has an ion intensity similar to only moderately abundant endogenous LPE and Cer species and appear close to the noise limit in the Orbitrap data. The increased sensitivity for LPE and Cer could arise from an enhanced transmission of lower *m/z* species under the employed settings or may reflect an enhanced ionisation resulting from the different ion source designs. The high concentrations reported for Cer 42:2;O2 likely arise due to some in-source fragmentation of the abundant HexCer 42:2;2 which also contributes to the ion signals. Using the HexCer IS we found the timsTOF data resulted in approximately 6% fragmentation of HexCer to Cer, compared to only ~1.5 % using the Orbitrap. This highlights the importance of considering in-source fragmentation effects when interpreting Q-MSI data and the utility of internal standard monitoring in-source fragmentation.

**Figure 4.**
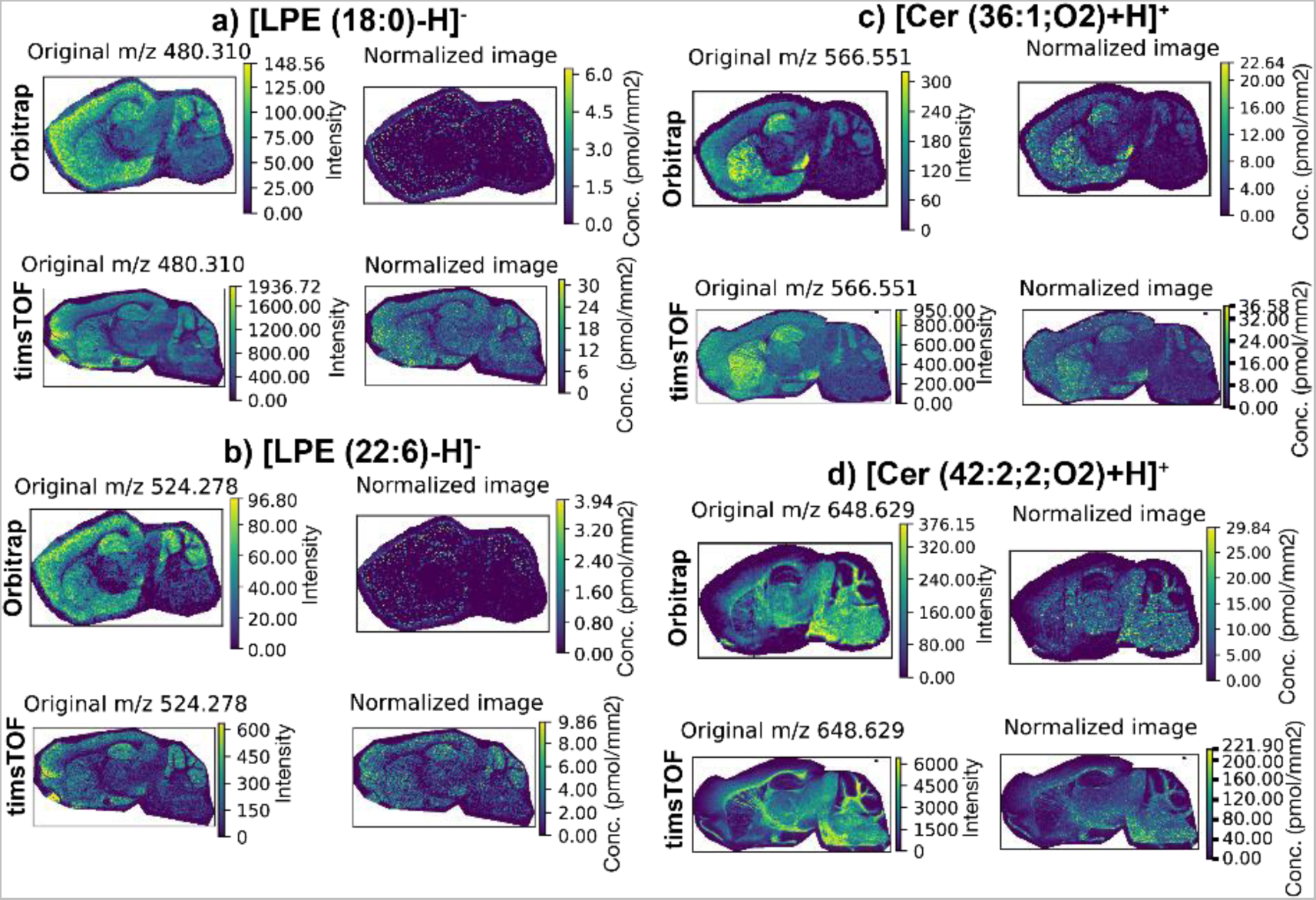
Comparison of Cer and LPE ion images obtained using the Orbitrap Elite and timsTOF systems; (a) [LPE(18:0)-H]^−^, (b)[LPE(22:6)-H]^−^, (c) [Cer(36:1;O2)+H]^+^ and d) Cer(42:2;O2)+H]^+^. The higher quality images obtained for the timsTOF arise due to better detection of the LPE and Cer IS peaks.

We next investigated the correlation in quantitative values obtained between the Orbitrap and timsTOF instrument by comparing the concentrations across whole tissue sections. **Figure 5** shows this correlation analysis for (a) SHexCer [M-H]^−^, (b) PC ([M+H/Na]^+^, (c) HexCer ([M+H]^+^, (d) PS, [M-H]^−^ (e) PE, [M-H]^−^ and (f) PI [M-H]^−^ lipid species. Correlation plots for additional lipid classes can be found *Supporting Information Figure S11*. These data also demonstrate the Q-MSI of lipids species covering almost 3 orders of magnitude. Encouragingly, high correlation of quantitative values (pmol/mm^2^) was obtained for most lipid classes that were well detected by both instruments. As expected, outliers were observed which were generally believed to be due to artificially elevated intensities in the timsTOF data given the lower mass resolution and, consequently isobaric interferences for some lipid masses. Many of these are attributed to the common type II isobaric overlap resulting from the [M+2] isotope of lipids containing one less double bond that are not resolved on the timsTOF but are resolved on the Orbitrap (e.g., [M-H]^−^ ions of PI 38:4[^13^C_2_] and PI 38:3). Here we have deliberately decided to show these isobaric overlaps to highlight limitations of limited mass resolution, however isotope correction could be performed to correct many of these outliers. Other outliers can be explained by an apparent increased sensitivity for certain lipid species on one instrument platform under the employed conditions. For example, several low abundance HexCer species such as HexCer 34:1;O2 and HexCer 44:1;O2 are observed at higher concentration using the timsTOF. This can be explained by the better detection of these HexCer species using the timsTOF. On the Orbitrap system these species have single pixel intensities very close to the noise level meaning they are not detected in many pixels and resulting in a possible underestimation of their concentration. Nonetheless, Figure 5 demonstrates that for lipid well detected by both systems without major isobaric interferences a strong agreement in Q-MSI data is obtained using both approaches, even when the IS was deposited onto tissues in different laboratories.

**Figure 5.**
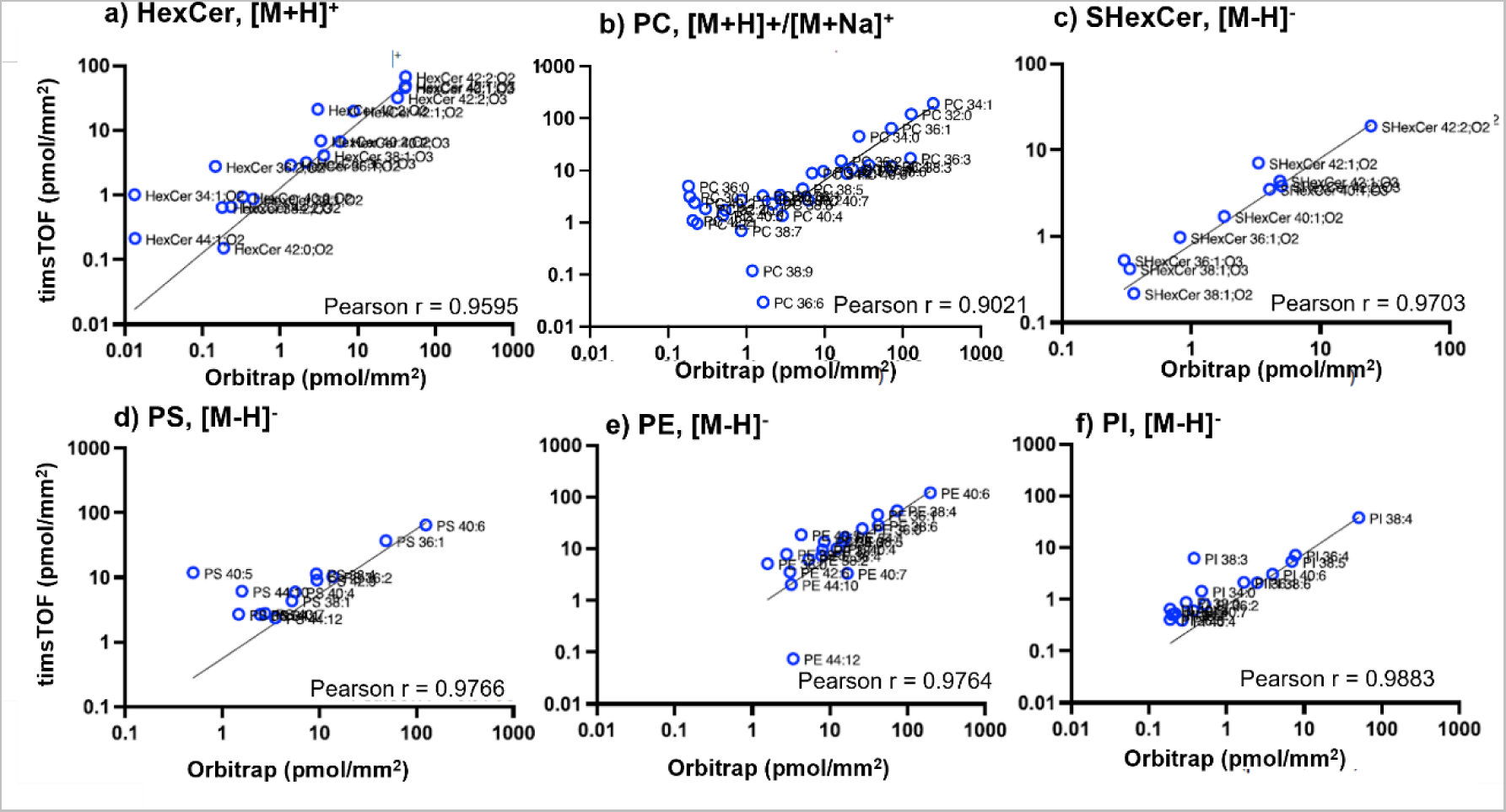
Correlation of Q-MSI data acquired using the timsTOF (mass resolution ~50,000 @*m/z* 750) and Orbitrap Elite (mass resolution ~180,000 @ *m/z* 750) following averaging of all on-tissue pixels for each section. Data is provided for (a) HexCer, (b) PC, (c) SHexCer, (d) PS, (e) PE and (f) PI lipid species. Each data point is the average of n=3 biological replicates measured on each system. The majority of outliers can be explained by isobaric overlap encountered in the lower resolution Q-TOF data which adds additional peak intensity in the extracted mass windows (see methods). In all cases P Correlation plots for additional lipid species are provided in *Supporting Information Figure S10*.

## Conclusion

By developing a multi-class IS mix and building on established quantitative workflows for shotgun lipidomics this study demonstrates an approach for the Q-MSI of lipids on an omics-wide scale (i.e., performing Q-MSI for multiple lipid species across multiple classes, simultaneously). The method can be readily adapted to other MSI modalities such as DESI, nano-DESI, IR-MALDESI and SIMS. Using the average concentrations across tissue sections, our data is consistent with quantitative amounts per mass of tissue reported in prior bulk lipidomics studies on mouse brain, demonstrating the validity of the approach. The use of MALDI-2 also facilitated Q-MSI of lipid species not typically well detected using conventional MALDI, such as the glycosphingolipids and Hex2Cer.

The robustness of our workflow is demonstrated by achieving similar Q-mSI results across two different laboratories and technical operators using both a higher mass resolution but lower throughput Orbitrap mass spectrometer and a much faster but lower mass resolution Q-TOF system. This multisite comparison yielded similar quantitative values for lipid species that are well resolved using both platforms and also provided insight into the number of lipid species for which quantitative errors may be encountered if an insufficient mass resolution is available to resolve isobaric interferences. Given the sensitivity of MALDI-2 signal to laser alignment the use of the IS mix is also shown to correct for MALDI-2-related artefacts that can lead to “striping “across ion images resulting in changing MALDI-2 ionisation efficiencies across the tissue (e.g., *Supporting Information Figure S4*). This approach can contribute to the robustness and comparison of MALDI-2 data acquired across long studies or across different laboratories. The approach can be further enhanced in the future by coupling with ion mobility methods to further remove possible isobaric interferences.

From a broader lipidomics perspective, this work provides an avenue to precisely determine lipidomes across tissue regions. With this, new metabolic information behind different pathologies such as neurodegenerative disease or cancer can be explored by revealing region-specific lipid compositional changes. It further creates an initial framework for the adaptation of lipid MSI to the established Lipidomics Minimal Reporting Checklist emphasizing on standardization and harmonization.^57^

## Associated Content

The supporting information is available free of charge at

Additional experimental methods and supplementary tables (Tables T1–T3) and figures (Figures S1–S11) as referred to in the main text.

## Acknowledgements

S. R. E acknowledges support from the Australian Research Council Future Fellowship Scheme (FT190100082). The authors are thankful for funding from the Michael J Fox Foundation (grant numbers MJFF-022753 and MJFF-019154). M. V and S. M. M contributed equally to this work.

## Conflicts of Interest

T. H. is an employee of Avanti Polar Lipids who provided the internal standards and have commercialised the IS Mix (MSI Splash®). N.V., M.C, N. H. P and M.J.Q.M. are employed by Aspect Analytics NV who performed the data analysis and commercialize software for MSI data analysis. N.V. and M.C. are shareholders of Aspect Analytics NV. K.E. is the owner of Lipidomics Consulting Ltd.

