## Supplementary Material for "Omics Scale Quantitative Mass Spectrometry Imaging of Lipids in Brain Tissue using a Multi-Class Internal Standard Mixture"

### Additional methods

### Haematoxylin and Eosin Staining

At UOW, post-MALDI imaged tissue sections were stained for histological and anatomical features by the haematoxylin and eosin (H&E) staining method. MALDI matrix was removed by immersion of slides in 100% methanol for 30 sec. Tissues were rehydrated by a series of graded ethanol washes of 95% ethanol (aq.) and 70% ethanol (aq.) and deionized water for 2 min each. Slides were stained by haematoxylin for 3 min, blued by rinsing in running tap water until clear and a 1 min wash in distilled water. Tissues were then stained in eosin for 30 sec, washed in two changes of 95% ethanol and once in 100% ethanol for 1 min each and transferred to xylene. Glass coverslip were placed onto samples by Quick hardening mounting medium. Digital optical scans of H&E-stained section were acquired at 10x magnification on a Falcon SP8 Confocal microscope (Leica Systems, Germany).

At UM, H&E staining was performed on the same 12-µm sections used for MALDI-MSI experiments. The residual matrix was removed by submerging the slides in 70% ethanol for 3 minutes. After a brief wash in Mill-Q water, the slide was re-submerged in 70% ethanol for 3 minutes, followed by Milli-Q water for 3 minutes. The slides were stained in hematoxylin for 3 minutes, followed by rinsing under running tap water for 3 minutes, then placed in eosin for 10 seconds. After placing the slides under running tap water for 1 minute, they were left in 100% ethanol for 1 minute followed by xylene for 30 seconds. The stained sections were mounted with Entellan and covered with a glass coverslip, then dried at room temperature. The stained sections were scanned with a digital scanner (Aperio CS2) at 20x magnification.

For both UM and UOW, high-resolution digital optical H&E images of tissue sections were uploaded onto a custom annotation portal for the annotation of key brain features namely prefrontal cortex, midbrain, hindbrain, basal ganglia and cerebellum. These features were coregistered to MSI data (.imzML) for extraction of regions of interest (ROI) for downstream analyses of lipid profile distributions.

**Table S1.** Internal standard concentrations. Spraying conditions: Flow rate = 0.06 mL/min, time = 1.55 min, surface area sprayed = 1,150 $mm^2$, number of layers = 16 passes, dilution factor = 0.10.

| Name | Molecular Formula | Average Mass (Da) | Concentration (μg/mL) | Total (μg/$mm^2$) | Picomoles/$mm^2$ |
|---|---|---|---|---|---|
| 15:0-18:1 (d7) PA (Na salt) | C36H61D7O8PNa | 689.927 | 111.00 | 0.144 | 20.82 |
| 15:0-18:1 (d7) PE | C38H67D7NO8P | 711.013 | 100.00 | 0.129 | 18.20 |
| 15:0-18:1 (d7) PG (Na salt) | C39H67D7O10PNa | 764.006 | 49.00 | 0.063 | 8.30 |
| 15:0-18:1 (d7) PI (NH4 salt) | C42H75D7O13P | 847.116 | 23.00 | 0.030 | 3.51 |
| 17:0-16:1 (d5) PS (Na salt) | C39H68D5NO10PNa | 775.000 | 105.00 | 0.136 | 17.53 |
| C12 Mono-Sulfo Galactosyl(β) Ceramide (d18:1/12:0) | C36H69N1O11S | 741.029 | 19.00 | 0.025 | 3.32 |
| 17:0 (d5) Lyso PE | C22H41D5NO7P | 472.610 | 3.00 | 0.004 | 0.82 |
| 15:0-18:1 (d7) PC | C41H73D7NO8P | 753.093 | 161.00 | 0.208 | 27.66 |
| 17:0 (d5) Lyso PC | C25H47D5NO7P | 514.700 | 3.00 | 0.004 | 0.75 |
| 15:0-18:1 (d7) PE | C38H67D7NO8P | 711.013 | 100.00 | 0.129 | 18.20 |
| 17:0 (d5) Lyso PE | C22H41D5NO7P | 472.610 | 3.00 | 0.004 | 0.82 |
| 18:1-18:1 SM (d9) | C41H72D9N2O6P | 738.140 | 31.00 | 0.040 | 5.43 |
| C15 Lactosyl(β) Ceramide (d18:1-d7/15:0) | C45H78D7NO13 | 855.210 | 13.00 | 0.017 | 1.97 |
| C18 Ceramide-d7 (d18:1/18:0) | C36H64D7NO3 | 572.997 | 11.00 | 0.014 | 2.48 |
| C17 Glucosyl(β) Ceramide (d18:1/17:0) | C41H79NO8 | 714.068 | 133.00 | 0.172 | 24.10 |

**Table S2.** Ions used for mass recalibration

EN- Endogenous Lipid, IS- Internal Standard

| Positive Ion Mode | | | | | |
|---|---|---|---|---|---|
| *Lipid* | *Type* | *Molecular Formula* | *Ion Adduct* | *m/z* | *Instrument* |
| PE 15:0-18:1(d7) | IS | C38H67D7NO8P | $[M+H]^+$ | 711.56642 | Elite |
| GluCer d18:1/17:0 | IS | C41H79NO8 | $[M+H]^+$ | 714.58784 | Elite , timsTOF |
| PC 15:0-18:1(d7) | IS | C41H73D7NO8P | $[M+H]^+$ | 753.61337 | Elite |
| PE 40:6 | EN | C45H78NO8P | $[M+H]^+$ | 792.55378 | Elite |
| **Negative Ion Mode** | | | | | |
| *Lipid* | *Type* | *Molecular Formula* | *Ion Adduct* | *m/z* | *Instrument* |
| SHexCer d18:1/12:0 ($NH_4$ salt) | IS | C36H72N2O11S | $[M-NH_4]^-$ | 722.45186 | Elite , timsTOF |
| PI 38:4 | EN | C47H83O13P | $[M-H]^-$ | 885.54985 | Elite |
| PI 15:0-18:1(d7) ($NH_4$ salt) | IS | C42H75D7NO13P | $[M-NH_4]^-$ | 828.56249 | Elite |
| SHexCer d42:2 | EN | C48H91NO11S | $[M-H]^-$ | 888.62401 | Elite |

**Table S3.** List of adducts used for Q-MSI of PC species. Adducts chosen to reduce isobaric interferences. The corresponding normalization IS mass is shown for each adduct type. ppm – parts per million.

| Theoretical | Measured | Formula | Species Level ID | Adduct | Normalization *m/z* | ppm Error |
|---|---|---|---|---|---|---|
| 706.5381 | 706.5381 | C38H76NO8P | PC 30:0 | [M+H]+ | 753.6134 | 0.00 |
| 730.5381 | 730.5381 | C40H76NO8P | PC 32:2 | [M+H]+ | 753.6134 | 0.00 |
| 732.5538 | 732.5538 | C40H78NO8P | PC 32:1 | [M+H]+ | 753.6134 | 0.00 |
| 734.5694 | 734.5694 | C40H80NO8P | PC 32:0 | [M+H]+ | 753.6134 | 0.00 |
| 758.5694 | 758.5695 | C42H80NO8P | PC 34:2 | [M+H]+ | 753.6134 | 0.13 |
| 760.5851 | 760.5851 | C42H82NO8P | PC 34:1 | [M+H]+ | 753.6134 | 0.00 |
| 762.6007 | 762.6004 | C42H84NO8P | PC 34:0 | [M+H]+ | 753.6134 | -0.39 |
| 786.6007 | 786.6007 | C44H84NO8P | PC 36:2 | [M+H]+ | 753.6134 | 0.00 |
| 788.6164 | 788.6163 | C44H86NO8P | PC 36:1 | [M+H]+ | 753.6134 | -0.13 |
| 790.6320 | 790.6322 | C44H88NO8P | PC 36:0 | [M+H]+ | 753.6134 | 0.25 |
| 800.5201 | 800.5202 | C44H76NO8P | PC 36:6 | [M+Na]+ | 775.5953 | 0.12 |
| 804.5514 | 804.5513 | C44H80NO8P | PC 36:4 | [M+Na]+ | 775.5953 | -0.12 |
| 806.5670 | 806.5689 | C44H82NO8P | PC 36:3 | [M+Na]+ | 775.5953 | 2.36 |
| 814.6321 | 814.6311 | C46H88NO8P | PC 38:2 | [M+H]+ | 753.6134 | -1.23 |
| 816.6477 | 816.6474 | C46H90NO8P | PC 38:1 | [M+H]+ | 753.6134 | -0.37 |
| 822.5044 | 822.5044 | C46H74NO8P | PC 38:9 | [M+Na]+ | 775.5953 | 0.00 |
| 824.5201 | 824.5197 | C46H76NO8P | PC 38:8 | [M+Na]+ | 775.5953 | -0.49 |
| 826.5357 | 826.5357 | C46H78NO8P | PC 38:7 | [M+Na]+ | 775.5953 | 0.00 |
| 828.5514 | 828.5512 | C46H80NO8P | PC 38:6 | [M+Na]+ | 775.5953 | -0.24 |
| 830.5670 | 830.5672 | C46H82NO8P | PC 38:5 | [M+Na]+ | 775.5953 | 0.24 |
| 832.5827 | 832.5830 | C46H84NO8P | PC 38:4 | [M+Na]+ | 775.5953 | 0.36 |
| 834.6007 | 834.6002 | C46H86NO8P | PC 38:3 | [M+Na]+ | 775.5953 | -0.60 |
| 842.6633 | 842.6626 | C48H92NO8P | PC 40:2 | [M+H]+ | 753.6134 | -0.83 |
| 844.6790 | 844.6782 | C48H94NO8P | PC 40:1 | [M+H]+ | 753.6134 | -0.95 |
| 852.5514 | 852.5497 | C48H80NO8P | PC 40:8 | [M+Na]+ | 775.5953 | -1.99 |
| 854.5670 | 854.5668 | C48H82NO8P | PC 40:7 | [M+Na]+ | 775.5953 | -0.23 |
| 856.5827 | 856.5824 | C48H84NO8P | PC 40:6 | [M+Na]+ | 775.5953 | -0.35 |
| 858.5983 | 858.5985 | C48H86NO8P | PC 40:5 | [M+Na]+ | 775.5953 | 0.23 |
| 860.6140 | 860.6141 | C48H88NO8P | PC 40:4 | [M+Na]+ | 775.5953 | 0.12 |
| 870.6946 | 870.6943 | C50H96NO8P | PC 42:2 | [M+H]+ | 753.6134 | -0.34 |
| 872.7103 | 872.7101 | C50H98NO8P | PC 42:1 | [M+H]+ | 753.6134 | -0.23 |

### Supporting Figure(s).

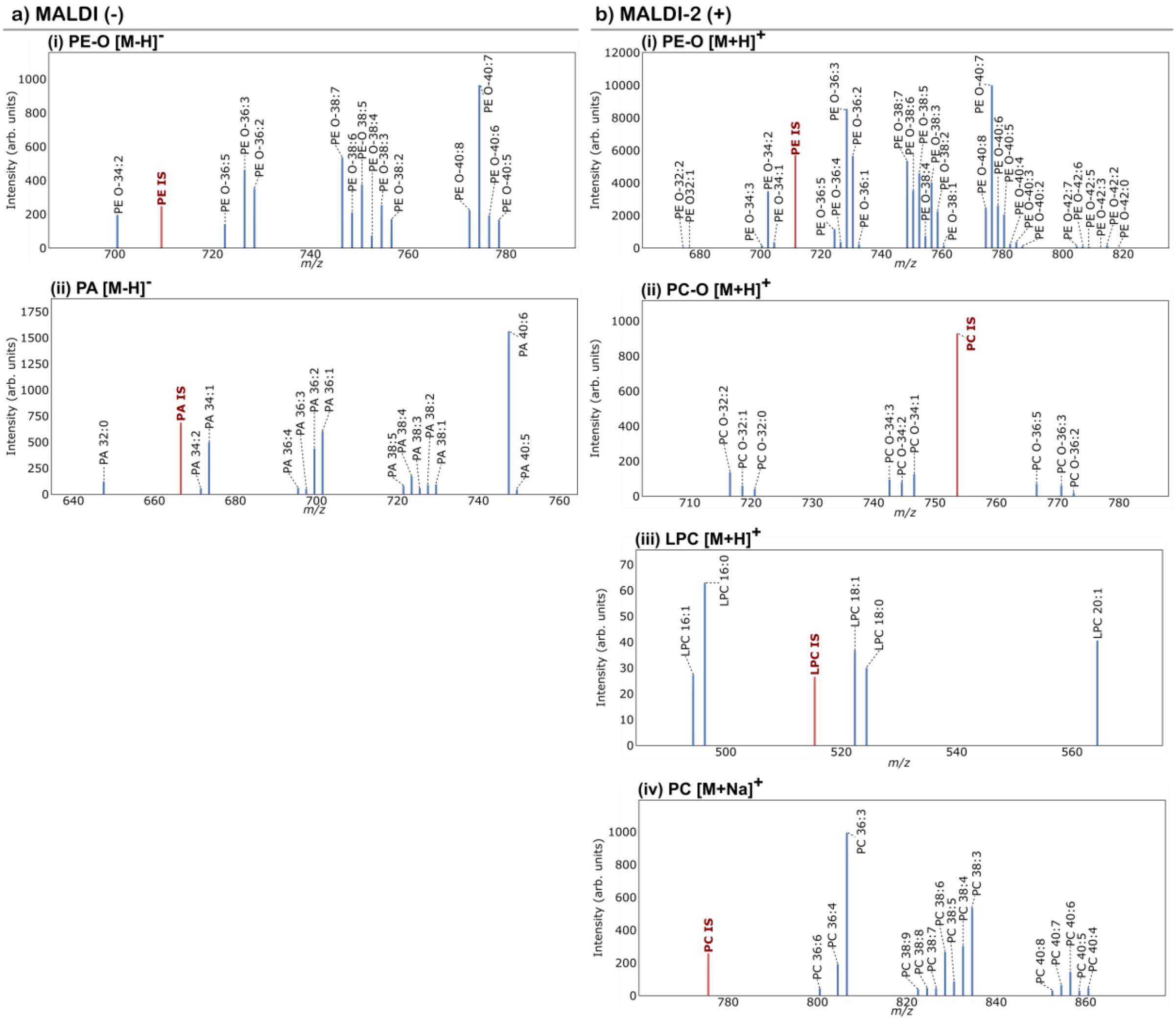

**Figure S1**. Extracted lipid species from positive ion mode analysis of mouse brain tissue. Reference IS peak shown in red and endogenous lipid species shown in blue; (a) Orbitrap Elite negative ion mode [M-H]$^-$ ions (i) PE-O and (ii) PA. (b) Orbitrap Elite positive ion mode: [M+H]$^+$ ions for (i) PE-O, (ii) PC-O, (iii) LPC and [M+Na]$^+$ ions for PC.

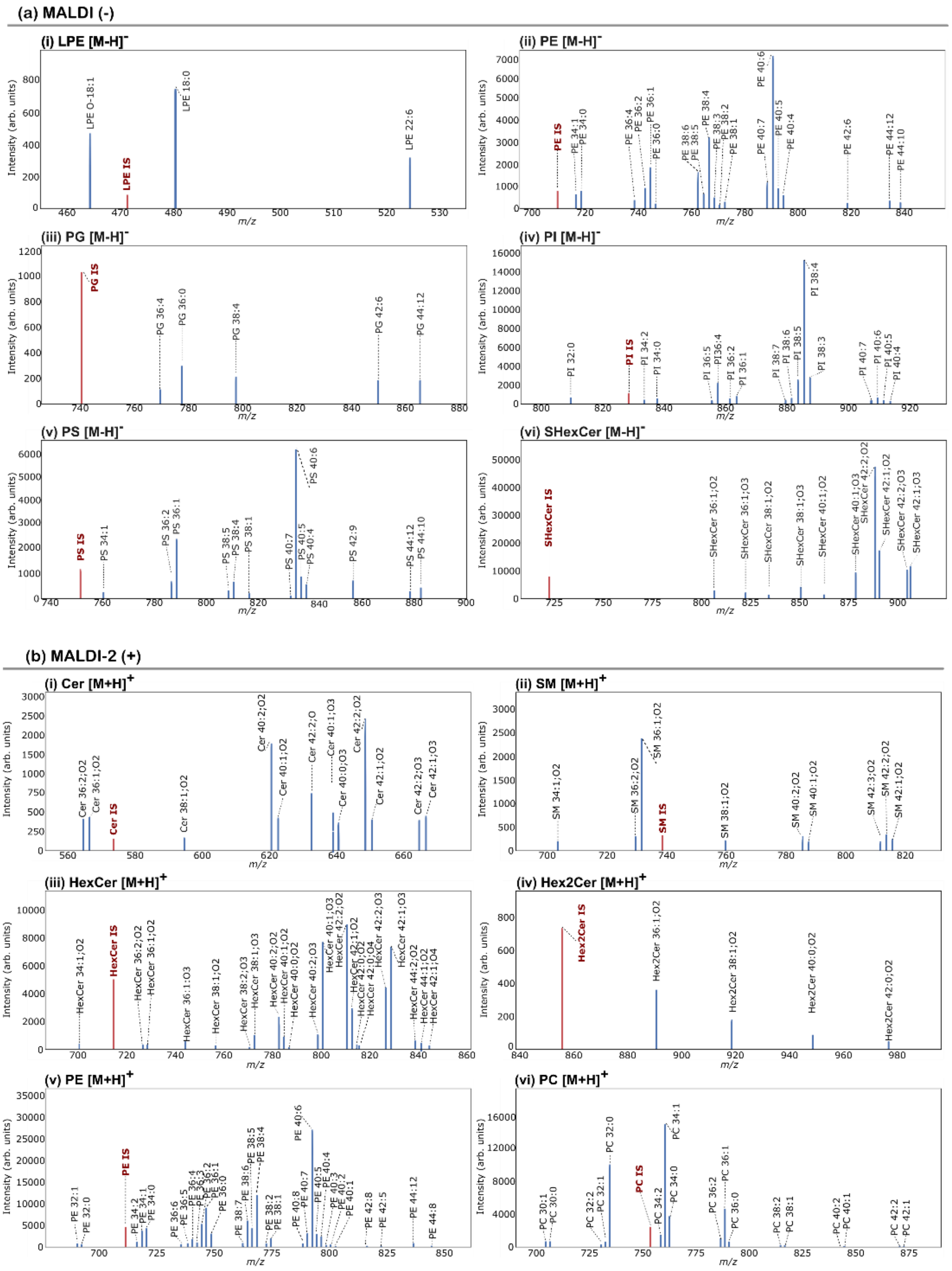

**Figure S2.** Extracted lipid species from negative ion mode analysis of mouse brain tissue. Reference internal standard (IS) peak shown in red and endogenous lipid species shown in blue; (a) timsTOF negative ion mode [M-H]$^-$ ions (i) LPE, (ii) PE (iii) PG, (iv) PI, (v) PS and (vi) SHexCer . (b) timsTOF positive ion mode: [M+H]$^+$ ions for (i) Cer, (ii) SM (iii) HexCer, (iv) Hex2Cer, (v) PE and (vi) PC.

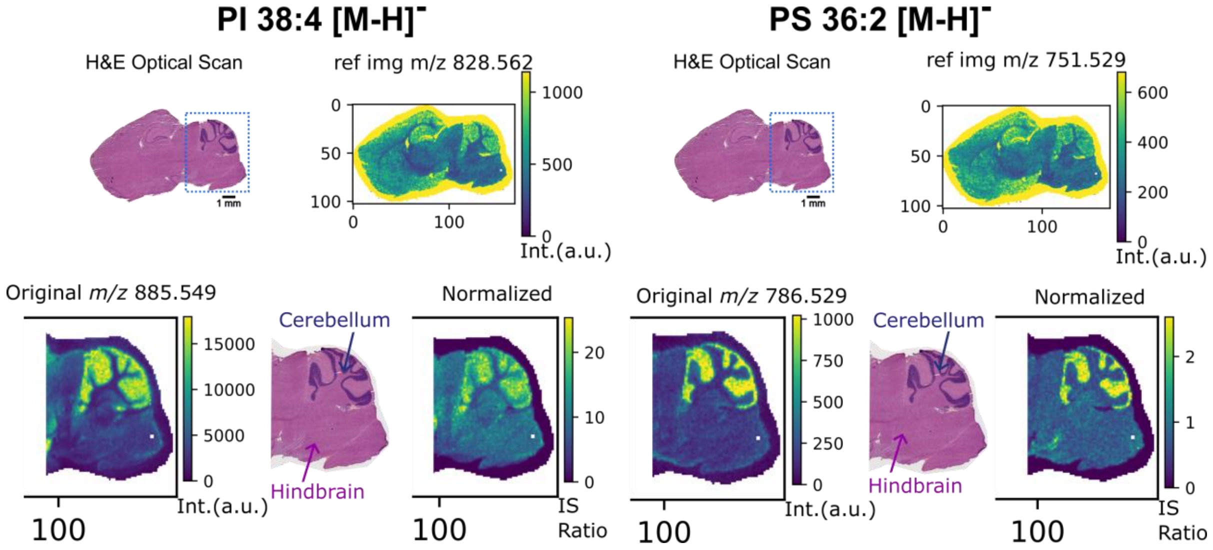

**Figure S3.** Benefit of internal standard normalization. Top panel; Left, [PI (38:4)-H]$^-$ *m/z* 885.5499 original ion image compared to IS normalized image reference *m/z* 828.5625. Right, [PS (36:42-H]$^-$ *m/z* 786.5291 original ion image compared to IS normalized image reference *m/z* 751.5291. Bottom panel; Blue outline region zoom into brain stem and cerebellum area depicting subtle increase in [PI 38:4 -H]$^-$ *m/z* 885.5499 and [PS 36:4 -H]$^-$ *m/z* 786.5291 within the hindbrain after IS normalization.

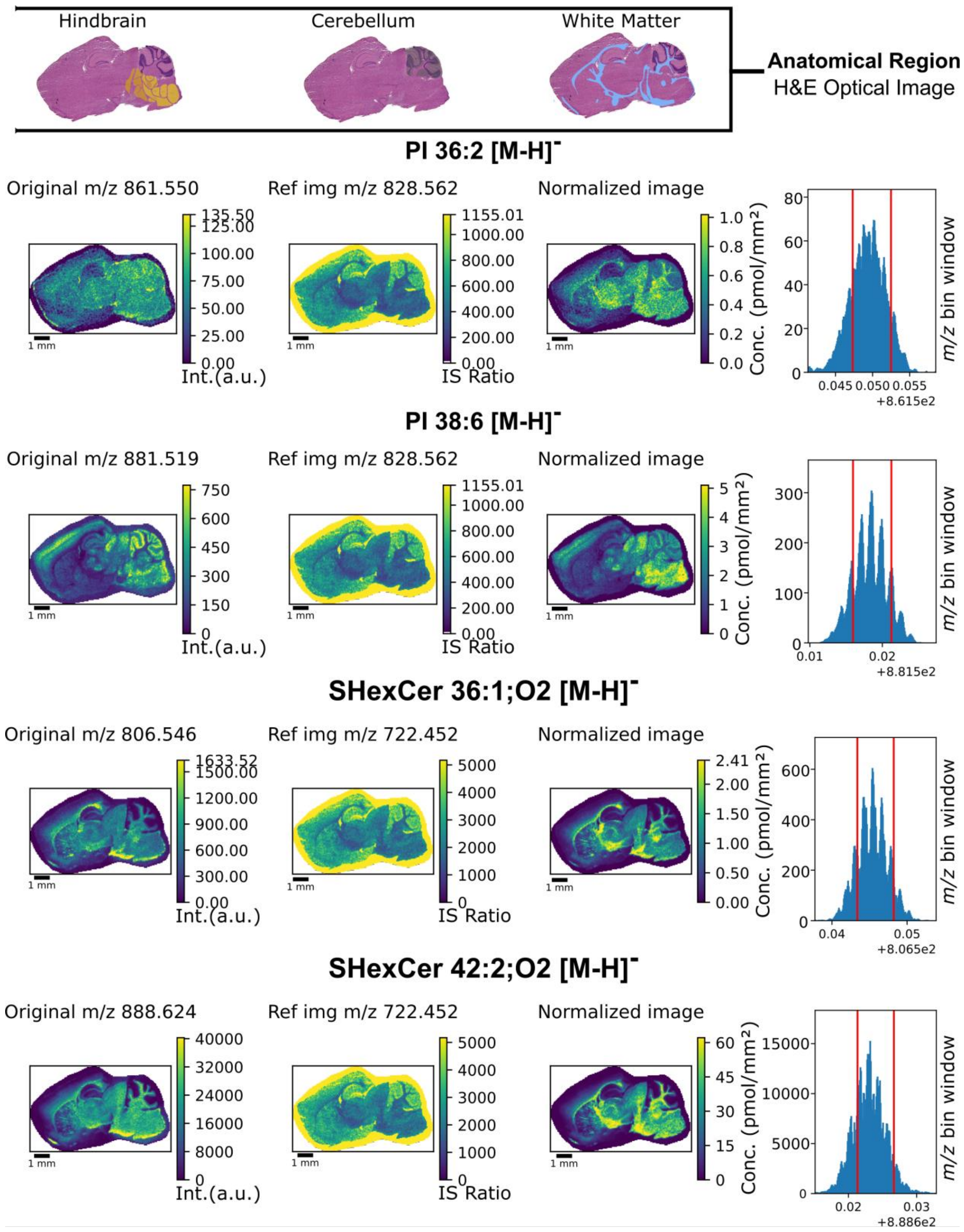

**Figure S4.** Selected Q-MSI data in negative ion mode. Quantitative mass spectrometry imaging of PI 36:2, PI 38:6, and SHexCer 36:1;O2 and SHexCer 38:1;O2 measured in negative ion mode. The right hand panel shows the 3 ppm selection window used for each species.

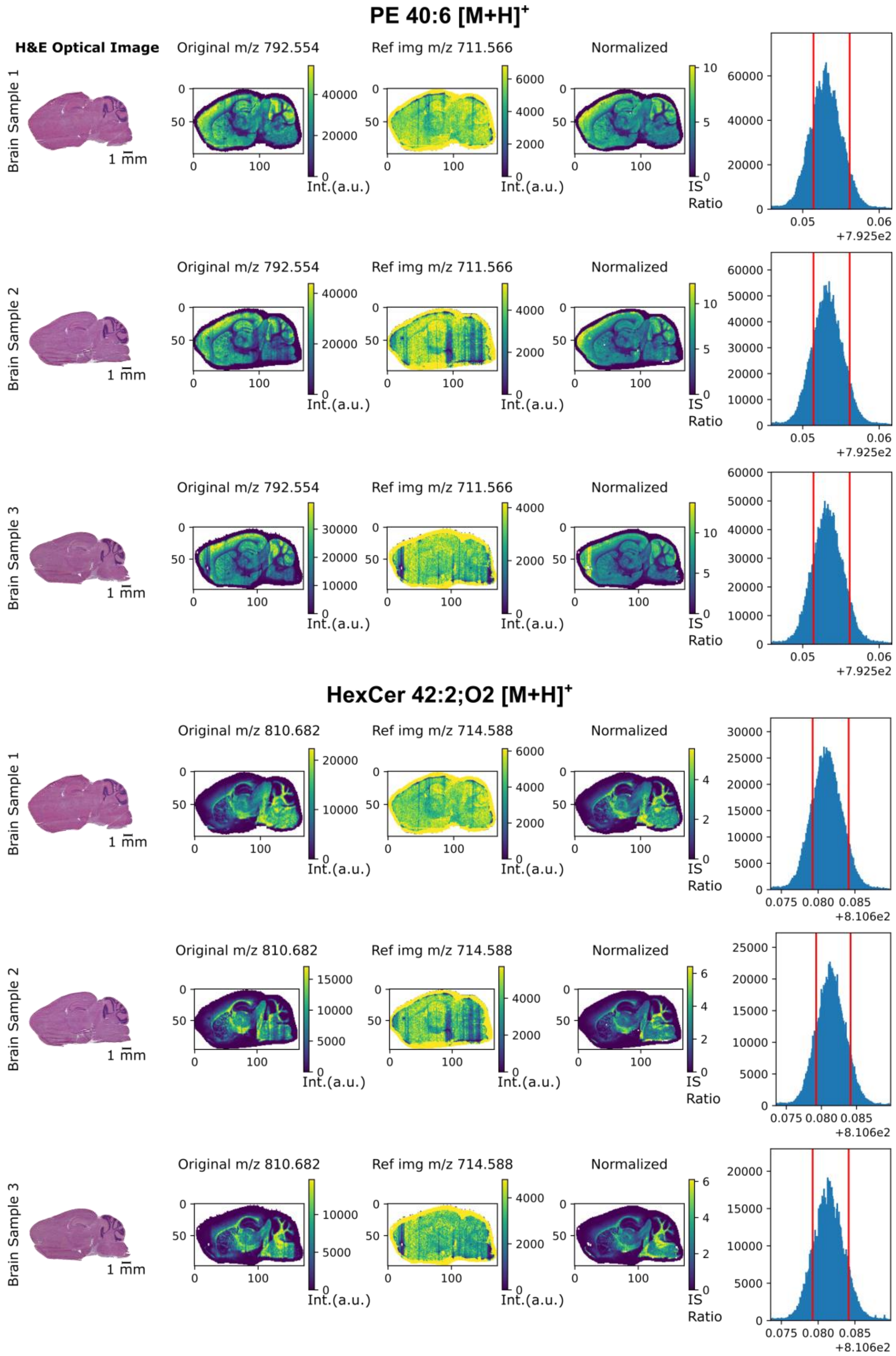

**Figure S5.** Reproducibility of quantitative mass spectrometry imaging. Figure shows PE 40:6 and myelin-rich HexCer 42:2;O2 species measured in positive-ion mode using MALDI-2. Data is generated from 3 biological replicates. IS normalized *m/z* images have significantly less stripes or line streaks in comparison to the original *m/z* images due to the correction of MALDI-2 artefacts (refer to main text). The right hand panel shows the 3 ppm selection window used for each species.

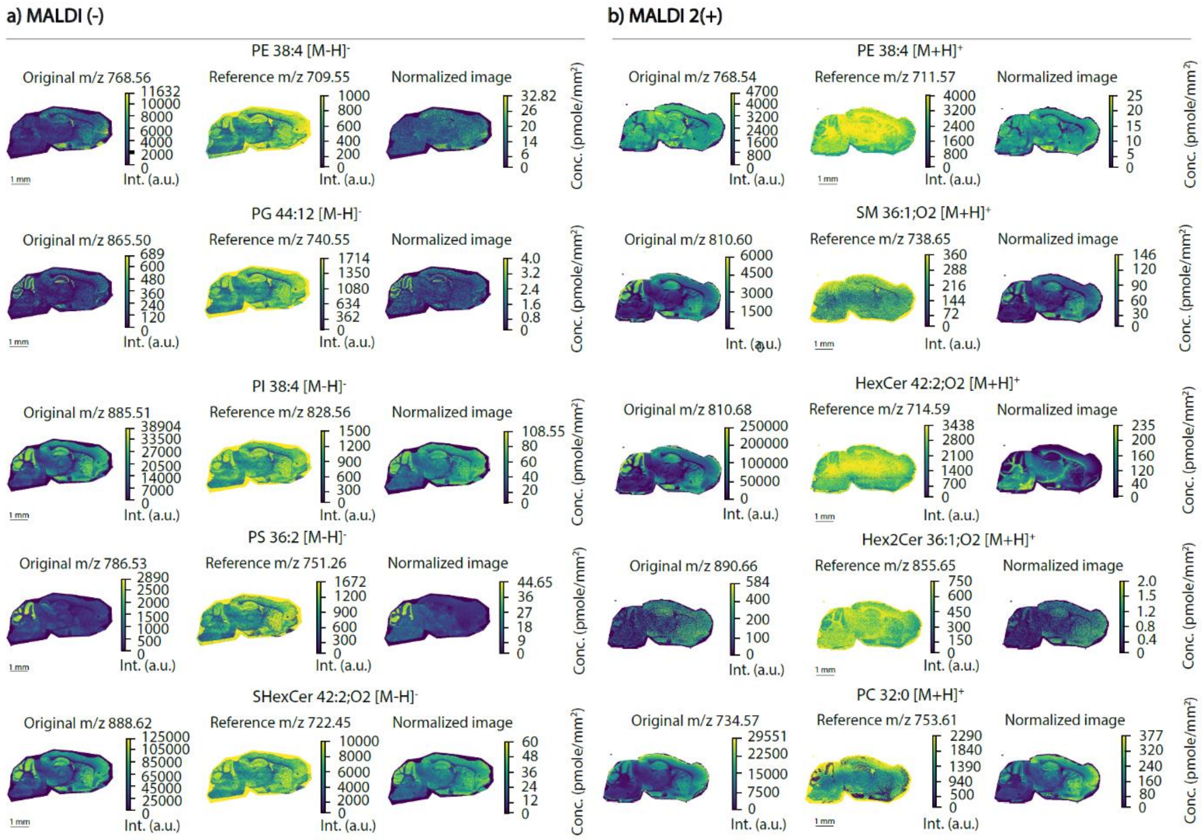

**Figure S6.** Representative internal standard normalized ion images acquired using the timsTOF. Figure shows different lipids species detected in (a) negative ion mode using MALDI and (b) positive ion mode using MALDI-2. For each lipid species the original ion image is shown on the left, the class-specific internal standard ion image is shown in the centre and the IS normalized ion image is shown on the right. Intensities for each lipid species were selected using an *m/z* window of ±12.0 ppm compared to the theoretical *m/z* of the lipid species (A) Left panel timsTOF MALDI-MSI negative ion mode detected as $[M\text{-}H]^-$; PE 38:4, PG 44:12, PI 38:4, PS 36:2 and SHexCer 42:2;O2. (B). Right panel timsTOF MALDI-2 MSI positive ion mode detected as $[M+H]^+$, PE 38:4, SM 36:1;O2, HexCer 42:2;O2, HexCer 42:2;O2 and PC 32:0.

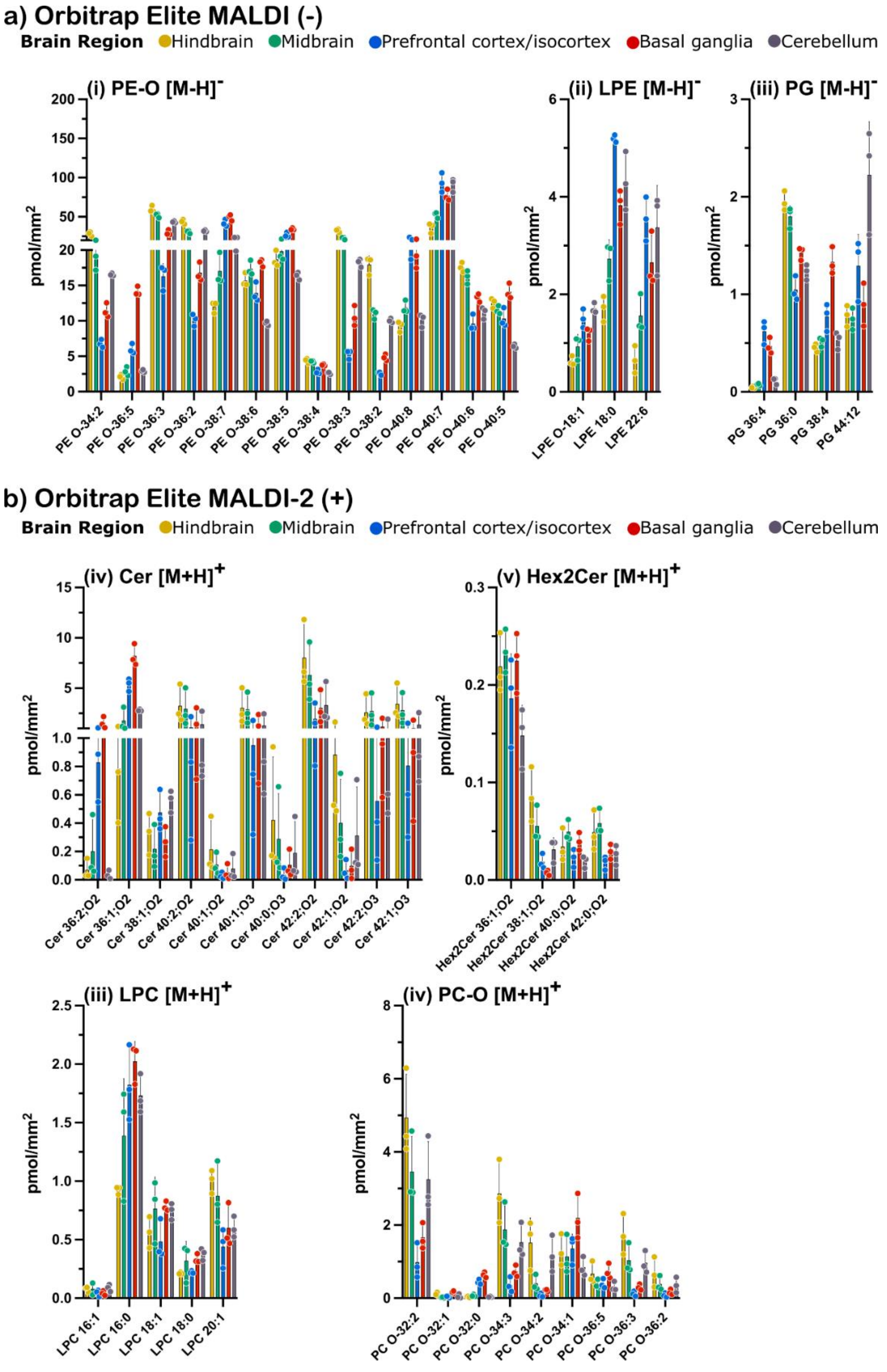

**Figure S7.** Region-specific mean lipid concentrations from Orbitrap Elite analyses. Regions of interest (ROI) from sagittal brain tissue sections are colour-coded hindbrain – orange, midbrain – green, prefrontal cortex/isocortex – blue, basal ganglia – red, and cerebellum – grey. (a) Quantitative MALDI-MSI negative mode $[M\text{-}H]^-$ ions for (i) PE-O, (ii) LPE/LPE-O and (iii) PG classes of lipids. (b) Quantitative MALDI-2 MSI positive mode $[M+H]^+$ ions for (i) Cer, (ii) Hex2Cer, (iii) LPC and (iv) PC-O species.

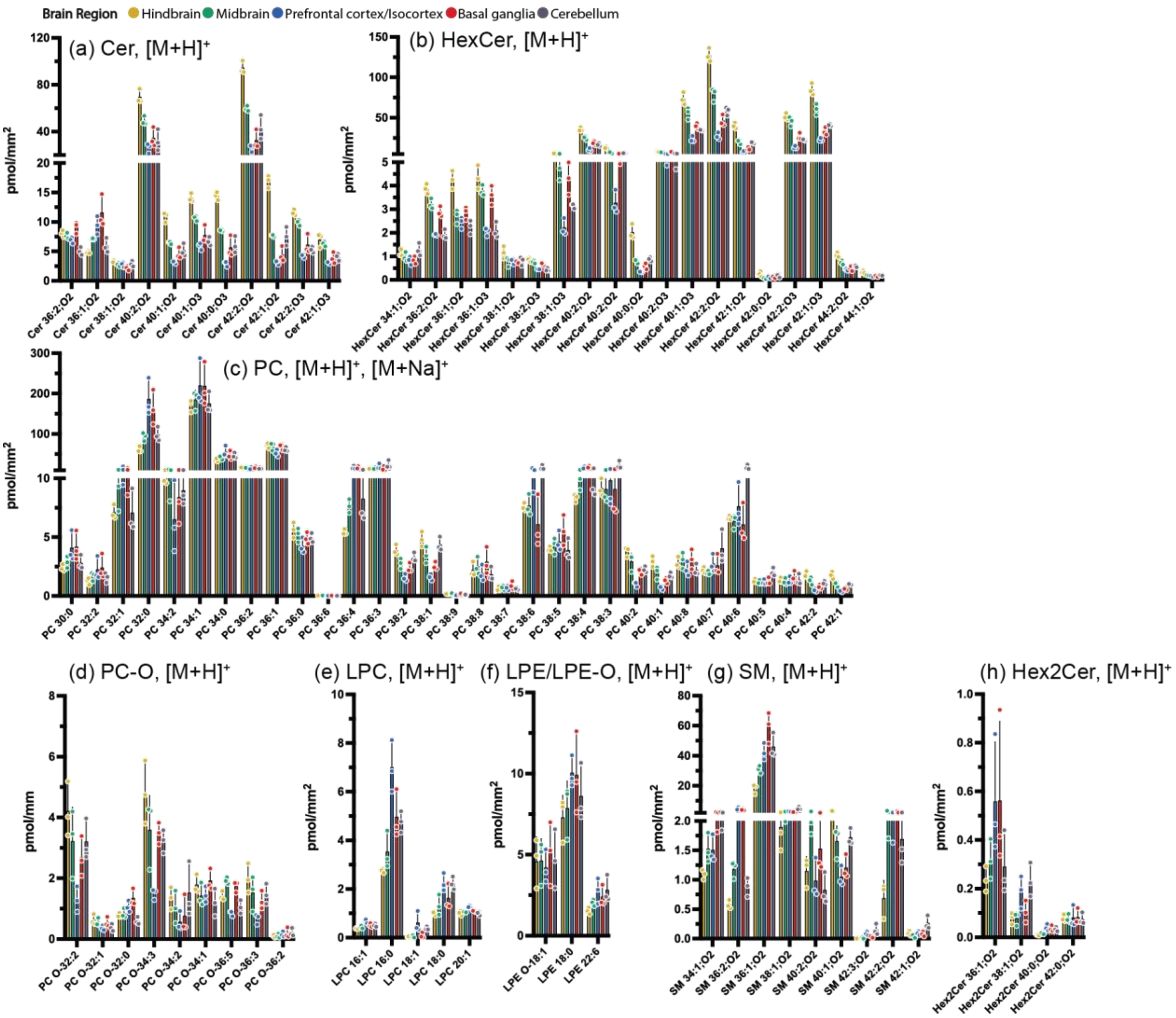

**Figure S8.** Region-specific mean lipid concentrations from timsTOF positive mode analyses. Regions of interest (ROI) from sagittal brain tissue sections are colour-coded hindbrain – orange, midbrain – green, prefrontal cortex/isocortex – blue, basal ganglia – red, and cerebellum – grey. Quantitative MALDI-2 MSI positive mode $[M+H]^+$ ions for (a) Cer, (b) HexCer, (c) PC, (d) PC-O, (e) SM, (f) LPC, (g) LPE/LPE-O species and (h) Hex2Cer species.

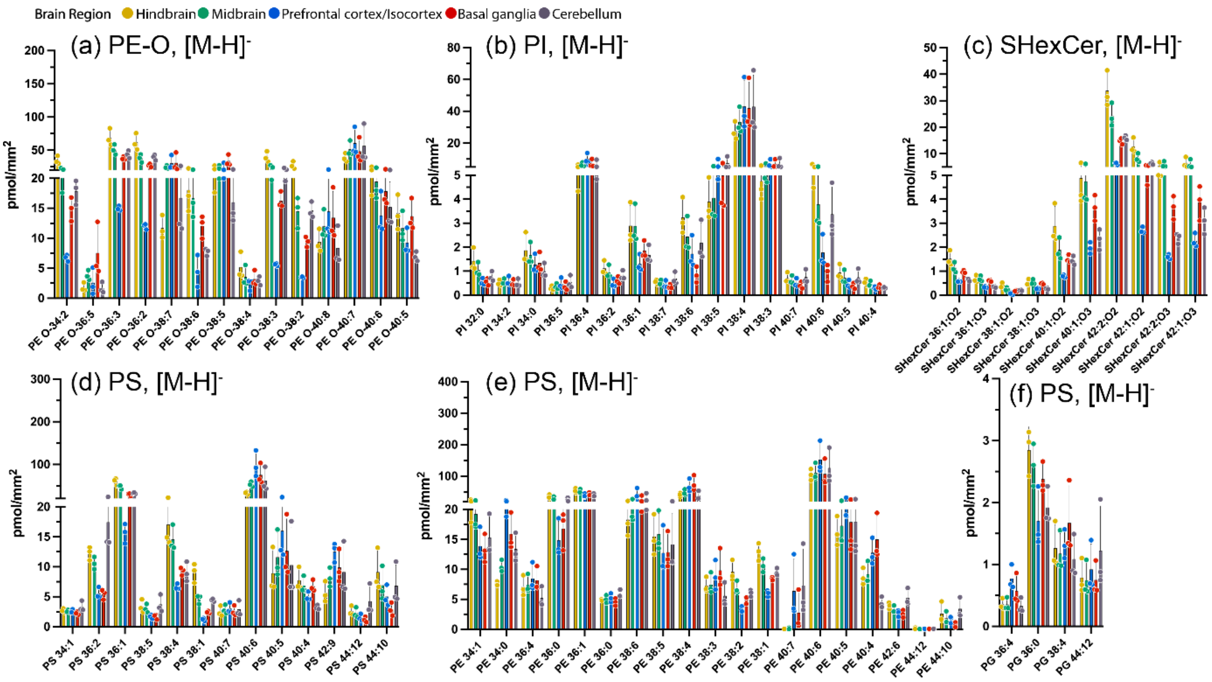

**Figure S9.** Region-specific mean lipid concentrations from timsTOF negative mode analyses. Regions of interest (ROI) from sagittal brain tissue sections are colour-coded hindbrain – orange, midbrain – green, prefrontal cortex/isocortex – blue, basal ganglia – red, and cerebellum – grey. Quantitative MALDI-MSI negative mode [M-H]- ions for (a) PE, (b) PG, (c) PE-O, (d), PI, (e) PS and (f) SHexCer classes of lipids.

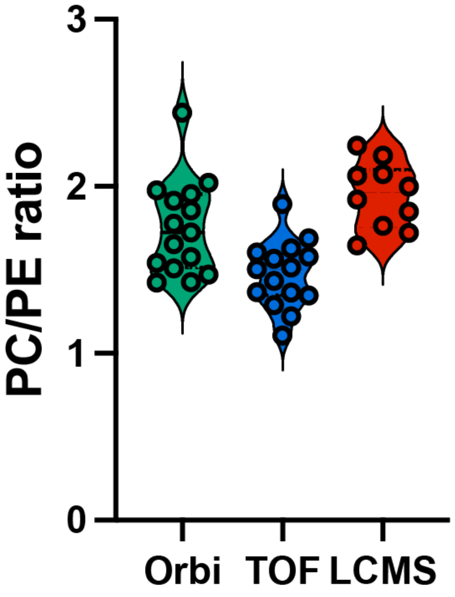

**Figure S10.** Violin plot comparing PC to PE ratio obtained by MALDI-MSI vs LC-MS/MS**.** Total PC and PE were obtained by summing up all class-specific species from respective analyses. PC to PE ratio was determined by dividing the total class levels. For MALDI-MSI, individual points represent a single measurement from a particular region, and for LC-MS/MS, measurements from extracted brain-homogenates of ten individual wild-type mice aged 24 weeks analyzed according to Zhang and colleagues (Zhang et al, *J. Lipid. Res*, 2022, 63, 100218 - doi: 10.1016/j.jlr.2022.100218)

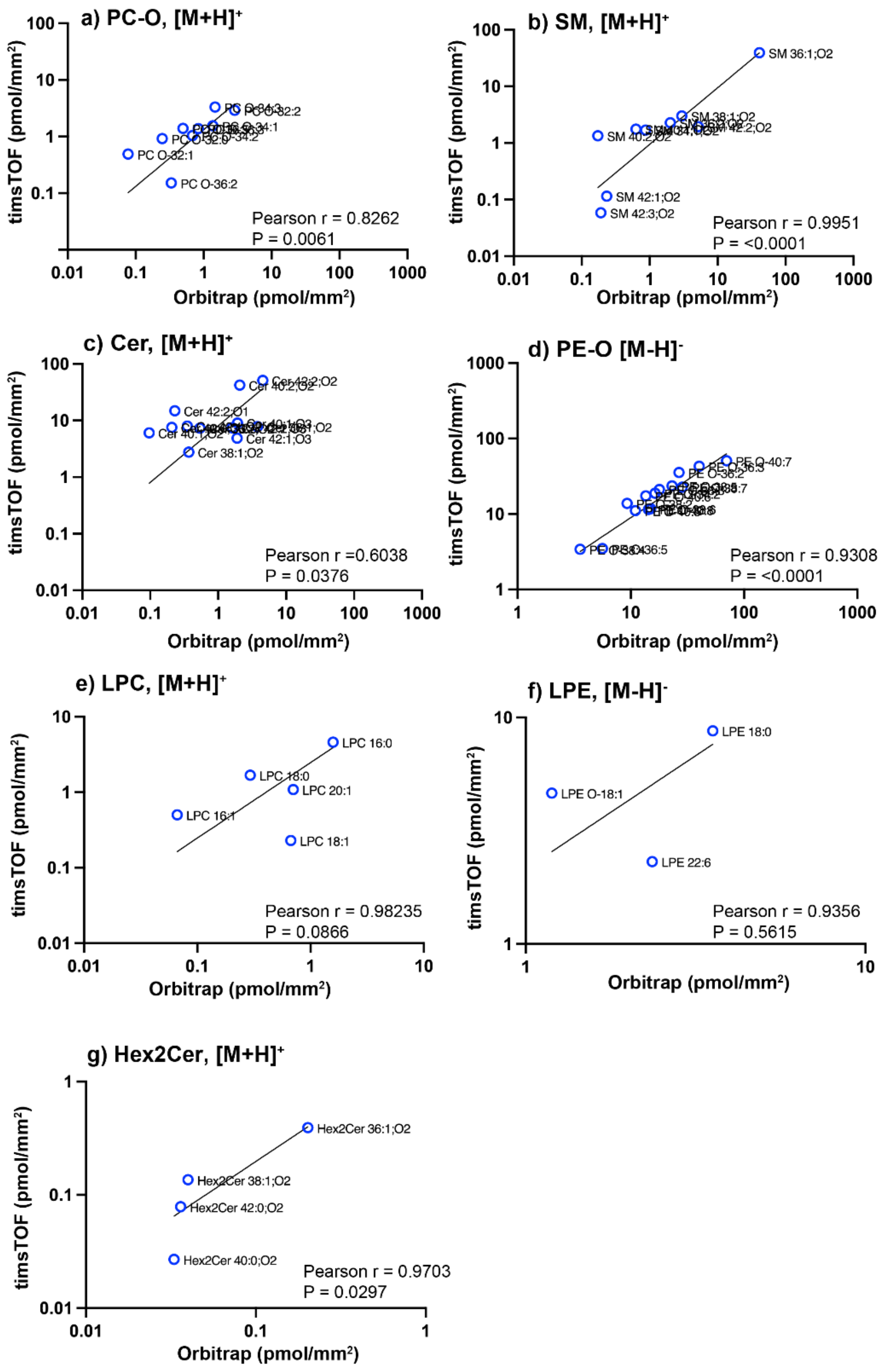

**Figure S11.** Correlation of Q-MSI Imaging data acquired using the timsTOF following averaging of all on-tissue pixels for each section. Data is provided for (a) PC-O, $[M+H]^+$, (b) SM, $[M+H]^+$ (c) Cer, $[M+H]^+$ and (d) PE-O, $[M-H]^-$, (e) LPC, $[M+H]^+$, (f) LPC, $[M-H]^-$ and (g) Hex2Cer, $[M+H]^+$. Each data point is the average of n=3 biological replicates measured on each system. The majority of outliers can be explained by isobaric overlap encountered in the lower resolution Q-TOF data which adds additional peak intensity in the extracted mass windows (see methods), or by the increased sensitivity of one system for a given lipid class, particularly for species that appear at low intensity that is close to the noise level in one or both systems.
